# Large-scale allosteric switch in the 7SK RNA regulates transcription in response to growth and stress

**DOI:** 10.1101/2021.09.16.460563

**Authors:** Samuel W. Olson, Anne-Marie W. Turner, J. Winston Arney, Irfana Saleem, Chase A. Weidmann, David M. Margolis, Kevin M. Weeks, Anthony M. Mustoe

**Affiliations:** Department of Chemistry, University of North Carolina, Chapel Hill, NC 27599-3290; Department of Medicine, University of North Carolina, Chapel Hill, NC 27599; UNC HIV Cure Center, University of North Carolina, Chapel Hill, NC 27599; Verna and Marrs McClean Department of Biochemistry and Molecular Biology; Therapeutic Innovation Center (THINC), Baylor College of Medicine, Houston, TX, 77030; Department of Molecular and Human Genetics; Baylor College of Medicine, Houston, TX, 77030

## Abstract

7SK is a highly conserved non-coding RNA that regulates eukaryotic transcription by sequestering positive transcription elongation factor b (P-TEFb). 7SK regulatory function likely entails changes in RNA structure, but characterizing dynamic RNA-protein complexes in cells has remained an unsolved challenge. We describe a new chemical probing strategy (DANCE-MaP) that uses maximum likelihood deconvolution and probabilistic read assignment to define simultaneously (*i*) per-nucleotide reactivity profiles, (*ii*) direct base pairing interactions, and (*iii*) tertiary and higher-order interactions for each conformation of multi-state RNA structural ensembles, all from a single experiment. We show that human 7SK RNA, despite significant heterogeneity, intrinsically codes for a large-scale structural switch that couples dissolution of the P-TEFb binding site to structural remodeling at distal release factor binding sites. The 7SK structural equilibrium is regulated by cell type, shifts dynamically in response to cell growth and stress, and can be exogenously targeted to modulate transcription in cells. Our data support that the 7SK structural ensemble functions as an integrator of diverse cellular signals to control transcription elongation in environment and cell specific ways, and establishes DANCE-MaP as a powerful strategy for comprehensively defining RNA structure and dynamics in cells.

## Introduction

RNA molecules fold back on themselves into complex secondary and tertiary structures that provide the basis of specific protein recognition, ligand binding, and broad gene regulatory functions (Sharp, 2009; Cech and Steitz, 2014). Most RNA elements can additionally fold into more than one structure, which can enable RNAs to function as regulatory switches (Dethoff et al., 2012). mRNA-based switches can regulate transcription, splicing, and translation of specific genes in response to metabolites (riboswitches) (Breaker, 2012) and protein binding (Ray et al., 2009; Fu et al., 2013). Large-scale RNA structural dynamics also underpin function of ribonucleoprotein (RNP) complexes such as the ribosome (Rodnina et al., 2017; Sengupta et al., 2019) and the spliceosome (Wilkinson et al., 2020). Nevertheless, despite their broad importance, RNA switches remain exceedingly difficult to identify, quantify in terms of their structure and in-cell equilibria, or link to functional outcomes.

The 7SK RNA is an abundant 332 nucleotide long non-coding RNA, forms the key architectural component of the 7SK small non-coding RNA-protein complex (snRNP), and serves as a major nexus of transcriptional control (Peterlin et al., 2012; Quaresma et al., 2016). 7SK is canonically thought to function by sequestering and inhibiting Cdk9/Cyclin T1 (together termed positive transcription elongation factor b, P-TEFb), a kinase required for phosphorylation and release of RNA polymerase II (Pol II) complexes paused at promoter-proximal regions. The P-TEFb-free form of 7SK appears to play additional roles in facilitating productive elongation (Peterlin et al., 2012; Quaresma et al., 2016), including modulating splicing (Barboric et al., 2009; Egloff et al., 2017; Ji et al., 2021), and chromatin remodeling (Eilebrecht et al., 2011; Flynn et al., 2016). The diverse functions of the 7SK snRNP are driven by coordinated changes in its bound protein components (Krueger et al., 2010). Methylphosphate capping enzyme (MePCE) (Yang et al., 2019) and La related protein 7 (LaRP7) (Krueger et al., 2008; Eichhorn et al., 2018) stabilize the core of the 7SK snRNP. P-TEFb is sequestered through interactions with the accessory protein dimer hexamethylene bis-acetamide inducible protein 1 or 2 (HEXIM1/2), which binds to the 7SK RNA at a high-affinity stem-loop structure, SL1 (Peterlin and Price, 2006; Czudnochowski et al., 2010; Martinez-Zapien et al., 2016). Under transcription stimulatory conditions, P-TEFb and HEXIM1/2 are liberated from 7SK by diverse release factors, including the bromodomain protein BRD4, and several helicases (Peterlin et al., 2012; Quaresma et al., 2016). The P-TEFb-free form of 7SK is in turn bound by heterogenous ribonucleoproteins (hnRNPs) and other proteins that interact with sites at the 3’ end the 7SK RNA (Peterlin et al., 2012; Quaresma et al., 2016). Most steps of this remodeling process remain poorly understood. Defining 7SK regulatory mechanisms will both illuminate fundamental aspects of transcriptional control and also inform ongoing efforts to inhibit transcription in disease settings, especially cancer (Olson et al., 2018), and, conversely, to activate transcription as part of “kick-and-kill” HIV cure strategies (Richman et al., 2009; Cary et al., 2016).

The 7SK RNA is highly structured (Wassarman and Steitz, 1991) and several studies support the model that P-TEFb binding and release involves remodeling of 7SK RNA structure (Krueger et al., 2010; Brogie and Price, 2017), or that the 7SK RNA exists in distinct conformations depending which proteins are bound (Krueger et al., 2010; Flynn et al., 2016; Brogie and Price, 2017). Multiple models for the 7SK RNA structure have been proposed (Wassarman and Steitz, 1991; Marz et al., 2009; Brogie and Price, 2017; Luo et al., 2021), but the accuracy of these models, whether they represent distinct co-existing states, and how these states might differentially modulate function remains unknown. To date, 7SK structure has primarily been studied using ensemble-average chemical probing approaches that are poorly suited for identifying coexisting RNA conformations or resolving in-cell structural dynamics. Typical of most non-coding RNAs, 7SK sequences show weak sequence covariation, precluding informative evolutionary analysis (Rivas et al., 2016; Kalvari et al., 2021). The 7SK RNP thus encapsulates broad features illustrating how RNA structural complexity endows functional complexity, and how such structural complexity frustrates mechanistic understanding.

Single-molecule chemical probing is emerging as a transformative technology for characterizing RNA structure and dynamics in living cells. The foundational conceptual advance is mutational profiling (MaP) reverse transcription, whereby a polymerase reads through and measures multiple chemical adducts per RNA molecule, recording them as mutations in complementary DNA (Homan et al., 2014). Massively parallel sequencing enables measurement of correlated modification events across hundreds of thousands of molecules, which encode rich information regarding RNA ensemble composition (Homan et al., 2014; Tomezsko et al., 2020; Morandi et al., 2021), and through-space secondary (Krokhotin et al., 2016; Cheng et al., 2017; Mustoe et al., 2019) and tertiary (Homan et al., 2014; Dethoff et al., 2018; Sengupta et al., 2019) structure interactions. However, existing single-molecule analysis frameworks only extract one type of information at a time (ensemble composition, base paring, or tertiary interactions). Ensemble deconvolution strategies permit measurement of multiple co-existing per-nucleotide reactivity profiles, but do not directly measure base pairs. Structures for each ensemble state can only be inferred, and this inference problem becomes increasingly ambiguous for long RNAs, particularly in cells. Conversely, the existence of multiple RNA structural states makes it challenging to assign base pairing and tertiary interactions.

Here we present a maximum likelihood (ML) strategy, DANCE-MaP (deconvolution and annotation of ribonucleic conformational ensembles) that extracts and annotates a large fraction of the total information from a single-molecule chemical probing experiment. DANCE-MaP directly visualizes complex RNA ensembles from MaP probing data, including direct detection of base pairs and tertiary interactions for each sub-state, at nucleotide resolution in a single experiment. We extensively benchmark this strategy in the adenine riboswitch, and our data reveal significant, previously undetected, complexity even within this well-defined RNA structural ensemble. We then apply DANCE-MaP to discover a large-scale, sequence-encoded structural switch in the 7SK RNA. Our structural-switch model rationalizes a large body of prior data and directly links P-TEFb release to concerted remodeling of 7SK structure in a cell-type and environmental specific way. We leverage this information to design a prototype, anti-sense oligonucleotide (ASO) to alter the 7SK ensemble and upregulate transcription in cells. Our work establishes DANCE-MaP as a powerful framework for directly resolving complex ensembles in cells and explains diverse features of 7SK lncRNA biology.

## Results

### Deconvolution of complex RNA ensembles with thermodynamic accuracy

RNA chemical probing data are conventionally analyzed by averaging across all molecules in a sample, yielding a single per-nucleotide reactivity profile that masks underlying dynamics (Fig. 1A, *bottom*). In single-molecule probing experiments, each MaP read represents a structural snapshot of an individual RNA molecule. For an RNA that folds into multiple structures, each structure yields distinct groups of reactive and unreactive nucleotides that are co-modified (or unmodified) in a correlated manner (Homan et al., 2014). We implemented a maximum likelihood (ML) framework that uses a modified Bernoulli mixture model to fit single-molecule reads to multiple reactivity profiles, sequentially increasing the number of fitted states until the optimal solution is identified (Fig. 1A, *right*; S1). Our ML strategy shares features with an independently described algorithm (Tomezsko et al., 2020), but additionally includes adjustments to handle missing data, improve robustness, and capture information from all four RNA nucleotides. Under idealized scenarios, our ML framework can deconvolute ensembles consisting of up to 5 states with populations ≥5% (Fig. S2). Deconvolution accuracy can break down as ensemble heterogeneity increases to ≥4 highly divergent states, meaning that individual deconvoluted states may still contain residual structural heterogeneity. Overall, reactivity profiles and populations are resolved with mean errors <1% for ensembles consisting of distinctive states.

**Figure 1:**
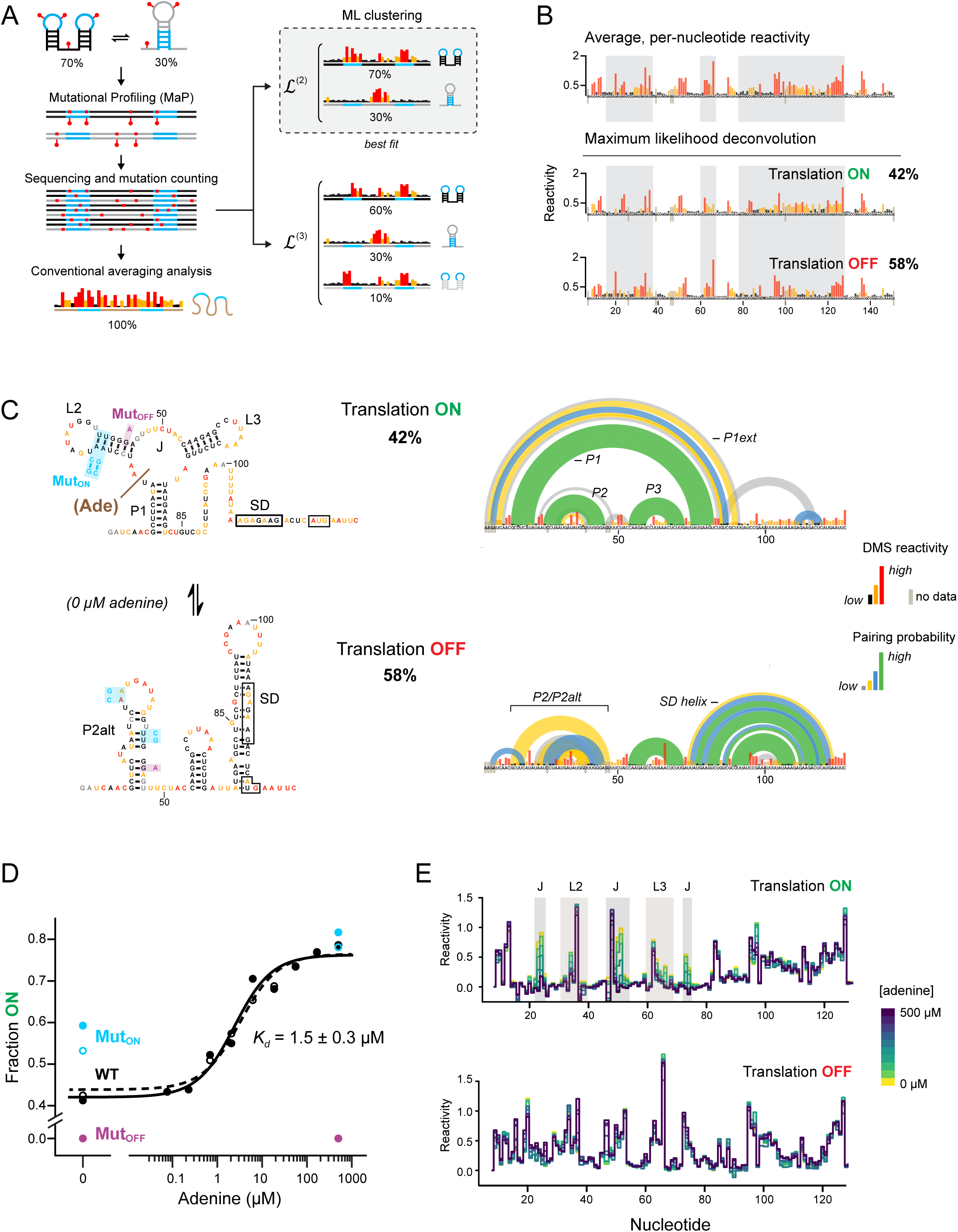
Maximum likelihood deconvolution enables thermodynamically rigorous analysis of RNA conformational ensembles. (A) Schematic of ML ensemble deconvolution. RNAs with multiple structures generate distinctive chemical modification patterns in single-molecule MaP data. For an RNA sampling multiple states, a (conventional) averaged, per-nucleotide reactivity profile may not be representative of any of the underlying structure states. ML analysis reveals both the individual reactivity profiles and populations of each ensemble state. (B) Averaged and deconvoluted DMS-MaP data for the ON and OFF states of the *add* adenine riboswitch. High, medium and low DMS reactivities are shown in red, yellow and black, respectively. Major reactivity differences between the ON and OFF states are shaded. (C) Deconvoluted structures of the *add* riboswitch. (*left*) Deconvoluted MaP data for the *add* riboswitch in the absence of ligand, superimposed on the NMR-defined (Reining et al., 2013) ON and OFF ensemble states. MutON (Reining et al., 2013) and MutOFF mutants are also illustrated. (*right*) State-specific pairing probabilities for the *add* ON and OFF states computed from deconvoluted reactivity profiles at 0 μM adenine. (D) Population of the ON state as a function of adenine concentration for native and mutant sequences. Replicates are shown in closed and open symbols. (E) Reactivities for the ON and OFF states as a function of adenine concentration. Regions that undergo adenine-dependent protection are emphasized with gray shading. Complete ensembles at each concentration are shown in Fig. S3 and S4. The *add* RNA is numbered per prior convention (Reining et al., 2013).

We validated our ML framework using the *V. vulnificus add* adenine riboswitch, which folds into a two-state ensemble consisting of a translation OFF state that occludes the Shine-Dalgarno (SD) sequence, and translation ON state that contains an adenine-binding aptamer (Fig. 1) (Reining et al., 2013). Structure probing experiments were performed on an *in vitro* transcribed RNA using dimethyl sulfate (DMS) under conditions where the reagent reacts with all four nucleotides (Mustoe et al., 2019). Whereas conventional averaging analysis suggests the riboswitch folds into a single well-defined structure, deconvolution of the single-molecule MaP data yielded two reactivity profiles matching the known translation ON and OFF states at the expected ∼40:60 populations (Fig. 1B) (Reining et al., 2013). Structure-specific differential reactivities are observed at both A and C residues and at U and G (for example, U84 and G85). These data enabled DMS-directed minimum free energy modeling and calculation of pairing probabilities that recapitulate the expected structures (Fig. 1C, *left*). Pairing probability analysis is a routine part of SHAPE-directed structure modeling and provides a key measure of model uncertainty (Siegfried et al., 2014; Smola et al., 2015a; Weeks, 2021), but until now has been unavailable for DMS data. Newly implemented here, pairing probability analysis reveals notable heterogeneity in the P2alt region in the OFF state, consistent with prior observations (Reining et al., 2013; Tomezsko et al., 2020), and with the potential for extended P1 pairing (P1ext) in the ON state (Fig. 1C, *right*).

We next examined the ability of ML deconvolution to measure biologically relevant changes in ensemble state by monitoring *add* riboswitch switching in response to adenine binding. Both OFF and ON conformations are populated even at saturating adenine ligand, consistent with prior studies (Warhaut et al., 2017; Tomezsko et al., 2020) (Fig. 1D, S3). The OFF state reactivity profiles are essentially identical at all adenine concentrations (Fig. 1E, *bottom*). By contrast, the ON state exhibits punctate reactivity changes within the ligand binding pocket, indicating adenine-induced stabilization of the aptamer domain tertiary structure (Fig. 1E, *top*). The concentration-dependence of these reactivity changes are consistent with the ML-resolved ON state representing a mixture of both ligand-free and ligand-bound states. Notably, the population of the ON state exactly follows the expected binding isotherm, enabling us to compute a Kd ≈ 2 μM in agreement with literature values (Reining et al., 2013) (Fig. 1D). The ability to resolve ligand binding thermodynamics with quantitative accuracy is unique to our study and provides rigorous validation of the precision of our ML deconvolution implementation. We further validated our approach using mutants that perturb the OFF–ON equilibrium. Consistent with other studies (Reining et al., 2013; Warhaut et al., 2017; Tian et al., 2018), destabilizing the P2alt helix via the MutON mutation only modestly shifts the equilibrium towards the ON state (Fig. 1E); our data reveal that this (unexpectedly) modest shift reflects creation of a new misfolded third state (Fig. S4). Destabilizing P2 via the MutOFF mutation (new to this study) switches the equilibrium to 65:35 between OFF and a new heterogenous state that is not adenine responsive (Fig. 1D, S4).

Collectively, these data validate the robustness and thermodynamic accuracy of our ML deconvolution framework, confirm that the *add* riboswitch functions via a conformational selection mechanism (Tian et al., 2018), and reveal unexpected complexity in the riboswitch sequence fitness landscape.

### Direct measurement of base pairs and tertiary interactions for individual RNA states

While ML deconvolution provides critical insight into ensemble composition and populations (Fig. 1, S2-S4), the set of RNA base pairs are not visualized directly and must be inferred via structure modeling. This inference problem is often ambiguous, especially for longer RNAs, where multiple sets of pairings can fit the same data, and for RNA structures measured in cells, where bound proteins complicate data interpretation (Smola et al., 2015b). Per-nucleotide reactivity data also do not report potential tertiary interactions. Single-molecule probing data contain characteristic correlations that directly measure through-space base pairing (PAIR, pairing ascertained from interacting RNA strands) and tertiary interactions (RING, RNA interaction groups) (Fig. 2A) (Homan et al., 2014; Mustoe et al., 2019). We developed a read assignment strategy which, in combination with our ML deconvolution approach, enables simultaneous measurement of per-nucleotide reactivity profiles and state-specific base pairing and tertiary interactions in complex structural ensembles. We term this integrated analysis framework <u>d</u>econvolution and <u>a</u>nnotation of ribo<u>n</u>ucleic <u>c</u>onformational <u>e</u>nsembles measured by <u>m</u>utational <u>p</u>rofiling, or DANCE-MaP.

**Figure 2:**
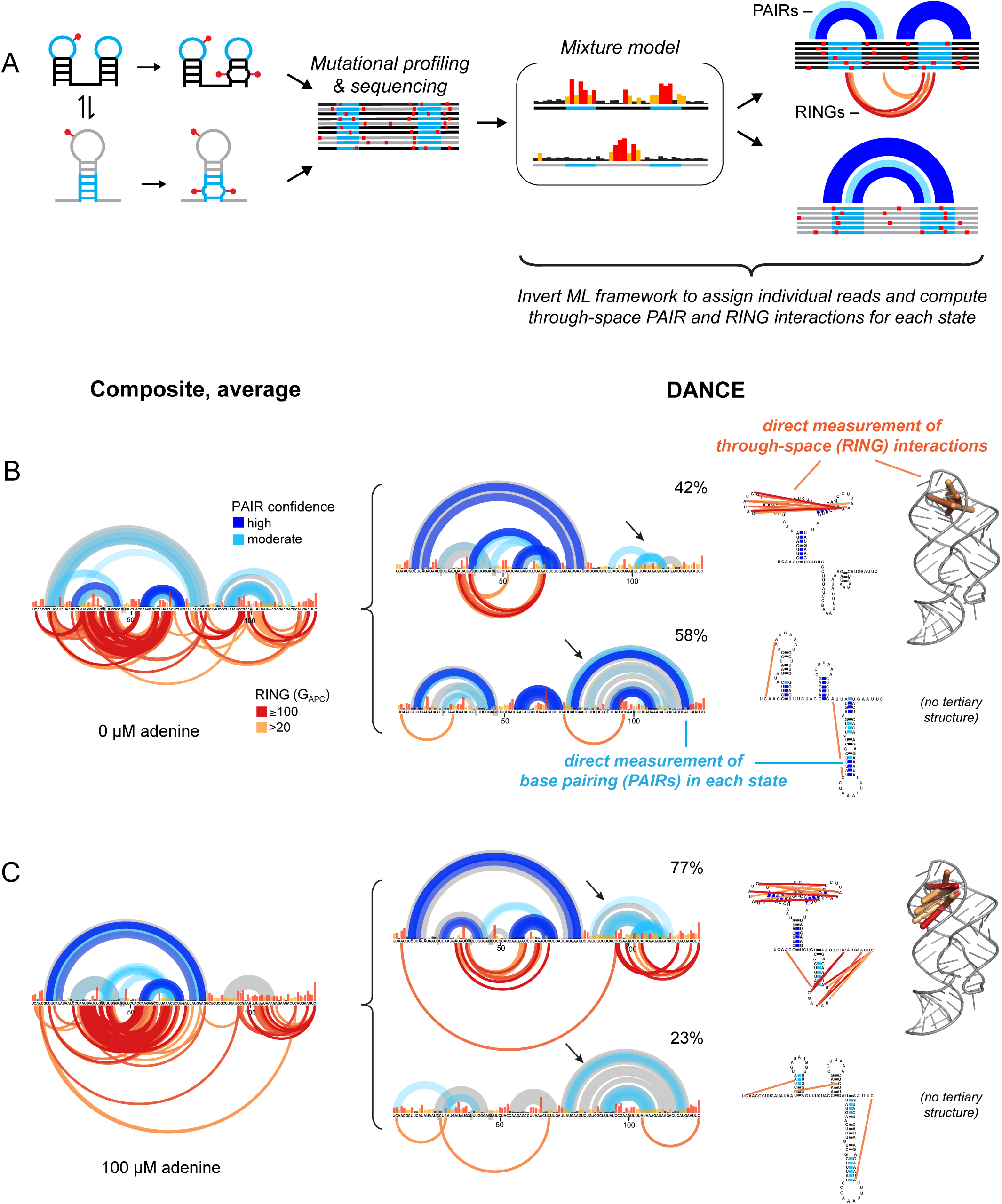
Direct DANCE-MaP detection of state-specific RNA base pairs and tertiary interactions. (A) DMS induces correlated disruptions of base pairing and tertiary interactions, measurable by MaP. Following ML deconvolution of per-nucleotide reactivity profiles, individual reads are assigned to distinct states. PAIR and RING correlation analyses are then used to directly detect base pairing and through-space tertiary interactions, respectively. (B,C) PAIR and RING analyses of composite (*left*) and deconvoluted (*right*) adenine riboswitch datasets, measured at 0 and 100 μM adenine. PAIRs are shown at top, superimposed on the modeled secondary structure state (light gray). High and moderate confidence PAIRs (denoted principal and minor in (Mustoe et al., 2019)) are dark and light blue, respectively. Arrows denote structure-specific PAIRs only observed in deconvoluted data. RINGs are shown at bottom, colored according to statistical significance. PAIR and RING correlations are also drawn on conventional secondary structure diagrams (*right*). RINGs measured in the ON state are superimposed on the crystal structure of the adenine riboswitch aptamer domain (far *right*; PDB: 4tzx (Zhang and Ferré-D’Amaré, 2014)).

Given a set of ML-deconvoluted reactivity profiles, individual single-molecule MaP reads can be assigned via a posterior probability calculation to their most likely parent state. State-specific PAIR and RING correlation analysis can then be performed on the assigned reads (Fig. 2A, S1). We devised a two-part strategy to measure state-specific PAIR and RING correlations and circumvent potential assignment biases (see Methods). Analysis of simulated MaP datasets confirmed that DANCE-MaP enables accurate measurement of state-specific RING and PAIR correlations (Fig. S5).

Without ML-deconvolution, PAIR and RING analyses of adenine riboswitch DMS-MaP datasets yield a dense meshwork of correlations consistent with the underlying structural dynamics, but which are challenging to interpret *de novo* (Fig. 2B, 2C, *left*). In contrast, DANCE-MaP reveals a specific and near-complete network of direct PAIR interactions that clearly define the ON and OFF state secondary structures (Fig. 2B, 2C, *right*; *in blue*). Remarkably, DANCE identifies base pairing interactions that are invisible in the composite data (Fig. 2B, 2C; *arrows*). Minor PAIR signals further reveal dynamics hidden by per-nucleotide analysis, including the known dynamics (Reining et al., 2013) of the P2/P2alt region in the OFF state. Equally striking, DANCE-MaP detected through-space tertiary interactions (RINGs) clearly identifying the L2-L3 loop-loop interaction in the ON state (Fig. 2B, 2C, *right*; *in red and orange*). Our data are consistent with other studies (Warhaut et al., 2017) showing that this tertiary interaction forms even in the absence of adenine ligand. Additional RINGs observed at the 3’ end of the ON state likely reflect further minor states. These data were reproducible across the comprehensive adenine titration (Fig. S3).

In sum, DANCE-MaP directly measures macrostate heterogeneity, base pairing, and tertiary interactions for each state in a complex ensemble, enabling complete structural analysis within a single, concise chemical probing experiment.

### Native 7SK RNA exists as a multi-state structural ensemble

Motivated by the fundamental role of 7SK RNA in transcriptional regulation and prior evidence of 7SK dynamics (Krueger et al., 2010; Flynn et al., 2016; Brogie and Price, 2017), we sought to define the 7SK structural ensemble and its role in regulating transcription. We performed DMS-MaP experiments on living human Jurkat cells and obtained high-coverage single-molecule DMS probing data for the 7SK RNA. Conventional averaged analysis (without deconvolution) yielded per-nucleotide reactivity profiles generally compatible with previously proposed SL1, SL3, and SL4 stem-loop structures (Wassarman and Steitz, 1991; Marz et al., 2009) (Fig. 3A, *top*). However, as observed previously (Wassarman and Steitz, 1991; Krueger et al., 2010; Brogie and Price, 2017; Wang et al., 2019), many nucleotides exhibit intermediate reactivities, consistent with significant, unresolved structural heterogeneity.

**Figure 3:**
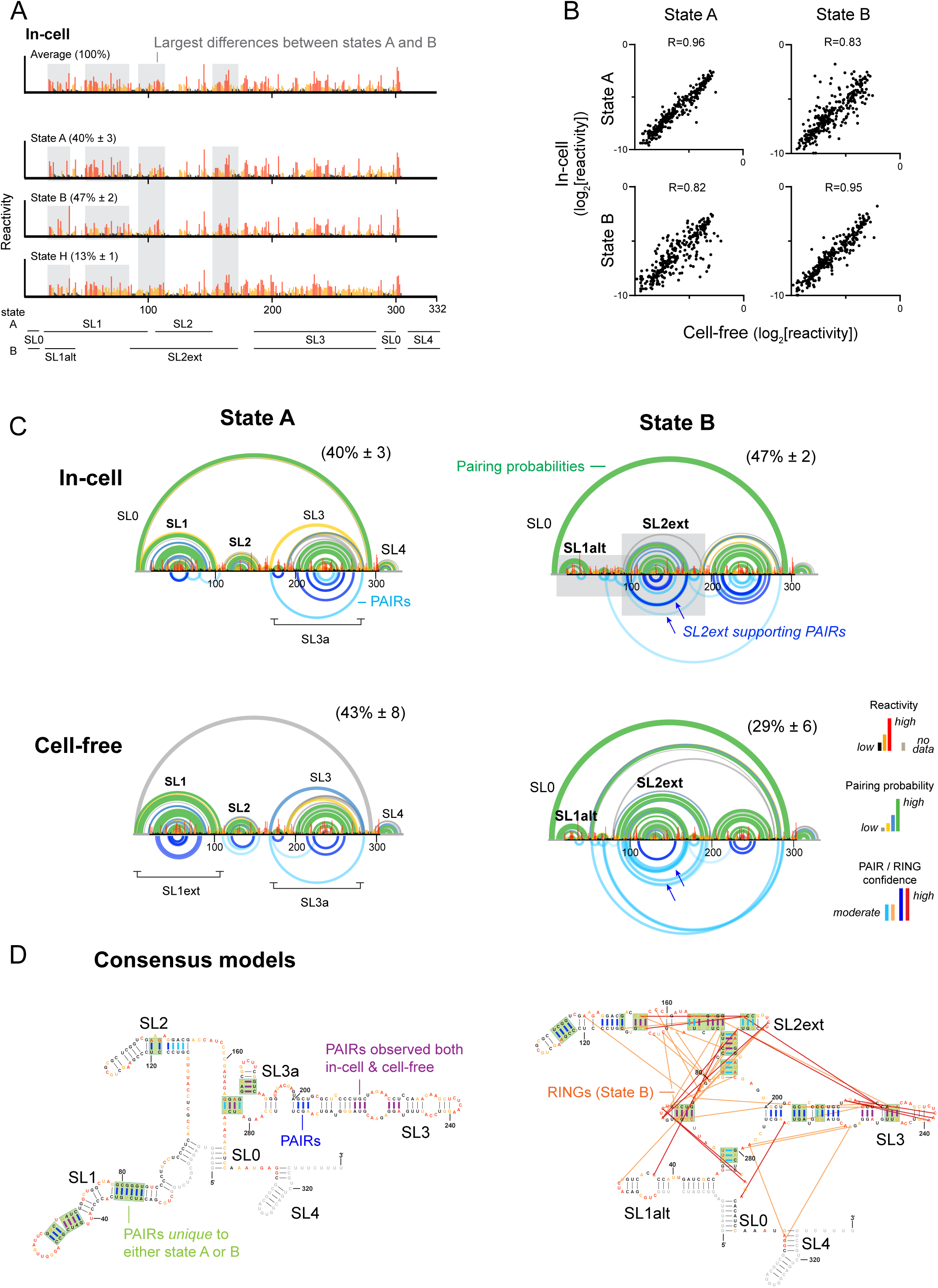
The 7SK RNA intrinsically codes for a large-scale structural switch. (A) Averaged and DANCE-deconvoluted reactivity profiles for 7SK RNA in cells. Major differences between states A and B are highlighted with gray shading. State H (heterogenous) has high reactivity throughout. Population averages and standard deviations are computed over 10 replicates (Table S2). Stem loop (SL) structural landmarks are indicated at bottom. (B) Comparison of state A and B per-nucleotide reactivities for in-cell and cell-free 7SK. Person’s R is shown. (C) In-cell and cell-free structural models for states A and B. Modeled base pairing probabilities (*top*) and directly measured PAIRs (*bottom*) are shown as arcs. (D) Consensus secondary structure models, shown in individual base pair format. Per-nucleotide reactivities are colored as per panel A. RINGs observed for state B are shown with orange-red lines, consistent with through-space structural communication and tertiary interactions. Measured PAIRs that directly support either state A or B are boxed in green; PAIRs observed in one versus both in-cell and cell-free datasets are shown with blue and magenta lines, respectively. Replicate data and state A and cell-free RINGs are shown in Figure S6.

ML single-molecule analysis indicated that 7SK structural heterogeneity reproducibly resolves into three states: A, B, and H (populations of 40% ± 3, 47% ± 2, 13% ± 2, respectively). State populations and reactivity profiles (R > 0.96) were highly reproducible over 10 biological replicates performed years apart. The minority H (heterogenous) state shares some features with B, but generally has high reactivity across the RNA. By comparison, the predominant A and B states show punctate regions of high and low nucleotide reactivity, consistent with these states representing well-defined structural states (Fig. 3A). Nucleotides throughout the SL1 region including U28, U30, U66, and U68 are unreactive in state A, but reactive in states B and H, corresponding precisely to nucleotides previously identified as changing conformation upon P-TEFb release (Krueger et al., 2010; Brogie and Price, 2017). Numerous additional differences occur throughout the 7SK RNA, indicative of a concerted global structural switch.

We repeated our experiments on protein-free RNA extracted from Jurkat cells that was heat denatured and refolded (referred to as cell-free). ML deconvolution revealed that cell-free 7SK also populates 3 states: A and B, and a mixed (M) state that shares features of both A and B (populations 43% ± 8, 29% ± 6, 28% ± 1; R > 0.98 between reactivity profiles; two consolidated replicates, see Methods). States A and B are the same as observed in-cell (R = 0.96 and R = 0.95 for states A and B, respectively Fig. 3B). Only diffuse reactivity protections and enhancements are observed relative to the in-cell RNA, which implies that the A and B states are dynamically rather than stably bound by proteins in cells. The lack of an H state under cell-free conditions is consistent with H representing a state heterogeneously bound by proteins in cells. Conversely, the lack of a mixed state in cells suggests that bound cellular factors specifically favor the A and B states. Thus, the 7SK RNA sequence intrinsically codes for two energetically balanced states, specifying a large-scale structural switch that behaves similarly with or without bound proteins.

### Direct base pair mapping and structure modeling reveals 7SK architecture

To fully resolve the secondary structure and potential tertiary structures of each state, we obtained high depth sequencing datasets (>3 million reads) that provide power sufficient to detect through-space PAIRs and RINGs across the 332 nt long 7SK RNA. These data reveal numerous PAIR signals that directly report base-paired structural elements distinctive to each state (Fig. 3C), reproducible between in-cell and cell-free environments, and across consolidated replicates (Fig. S6). We used these PAIR data in combination with per-nucleotide reactivity profiles to build detailed secondary structure models for the A and B states (Fig. 3D). The resulting structural models reveal that 7SK folds into two globally different conformations, each of which is supported by distinctive per-nucleotide reactivities and state-specific, direct PAIR correlations. Both states also show alternative predicted pairing possibilities and PAIRs suggestive of residual heterogeneity (Fig. 3C), indicating that states A and B should be interpreted as class averages rather than pure states. Nevertheless, each state possesses key defining structural features, and these state-specific structures clearly support a link between 7SK conformational dynamics to P-TEFb binding and release.

#### State A is the P-TEFb binding-competent state with a dynamic SL0 stem

State A largely recapitulates classic models of 7SK structure, blending features predicted by early probing studies (Wassarman and Steitz, 1991) and more recent evolutionary analyses (Fig. S7) (Marz et al., 2009). The SL1 helix is the defining structural feature of state A, and is directly supported by PAIRs both in cells and for the cell-free extracted RNA (Fig. 3C). SL1 has been extensively validated as the recognition site for HEXIM1/2 and P-TEFb, based on in vitro binding assays (Lebars et al., 2010; Martinez-Zapien et al., 2016), analysis of P-TEFb–bound 7SK fractions from cells (Brogie and Price, 2017), and in-cell functional assays (Egloff et al., 2006; Fujinaga et al., 2014). The population of state A in cells, 40%, is also consistent with the estimated fraction of 7SK bound by P-TEFb (Nguyen et al., 2001; Yang et al., 2001). Thus, we assign state A as the P-TEFb bound (sequestered) state.

Structure modeling indicates that this P-TEFb-sequestered state contains the long-range SL0 pairing interaction between the 5’ and 3’ ends that “circularizes” the RNA (Marz et al., 2009). While we lack data for the 5’ strand of SL0 due to overlap with the primer binding site, the 3’ strand of SL0 is lowly-to-moderately reactive in-cells, consistent with formation of a dynamic, partially stable stem (Fig. 3, S8A). By contrast, the alternative extended form of SL1, which out-competes SL0 in the cell-free RNA (SL1ext, see Fig. 3C), is reactive in cells, arguing against the “linear” structure. The increased stability of SL0 in cells likely reflects favorable interactions with MePCE and LARP7, bound at the 5’ and 3’ ends of the RNA, respectively (Muniz et al., 2013; Eichhorn et al., 2018; Yang et al., 2019); indeed, SL0 pairing facilitates MePCE-LARP7 interactions *in vitro* (Brogie and Price, 2017). Given that SL0 is strongly supported in state B (Fig. 3A, S8A), our data thus indicate that the 7SK RNP primarily exists in a “circular” form in cells.

Additional structural features include the SL2, SL3, and SL4 stems, proposed in prior studies (Wassarman and Steitz, 1991; Marz et al., 2009; Brogie and Price, 2017). PAIR signals provide the first direct validation of the SL2 and SL3 stems (Fig. 3C, 3D). PAIR analysis further reveals a long-range SL3a interaction present for both in-cell and cell-free RNAs (Fig. 3C, 3D, *left*). Finally, we performed RING analysis to search for potential tertiary interactions that may stabilize the A state, analogous to those validated for the adenine riboswitch (Fig. 2B, 2C). Observed RINGs are relatively isolated (Fig. S6), and thus we see no compelling evidence for tertiary interactions in state A.

#### State B is the P-TEFb released state with remodeled SL1 and central domains

State B constitutes a novel structure without close literature precedent (Fig. 3C, S7). Most notably, SL1 is absent. Instead, this region folds into the previously postulated SL1alt stem (Krueger et al., 2010; Brogie and Price, 2017). Although overlap with the primer binding site precludes measurement of SL1alt-specific PAIRs, the disappearance of the SL1 PAIRs (as observed in state A) implies that this region adopts an alternative structure in state B. SL1alt, and not SL1, is also clearly supported by per-nucleotide DMS reactivities and by pairing probabilities (Fig. 3C). P-TEFb does not bind SL1alt (Czudnochowski et al., 2010; Fujinaga et al., 2014) and, indeed, P-TEFb binding converts SL1alt to SL1 *in vitro* (Brogie and Price, 2017). Conversely, release of P-TEFb induces conversion of SL1 to SL1alt (Krueger et al., 2010; Brogie and Price, 2017). The 47% population of state B in cells is also consistent with the fraction of 7SK that is in a P-TEFb-released state in cells (Nguyen et al., 2001; Yang et al., 2001). Thus, we conclude that state B constitutes the P-TEFb released state.

SL1alt is coupled to formation of a major extension of SL2, which we term SL2ext, that has not been observed previously (Fig. 3C). Re-pairing to form SL2ext is directly supported by PAIRs in both in-cell and cell-free RNAs (Fig. 3C, blue arrows). Indeed, PAIR analysis was essential for resolving these interactions: SL2ext is not predicted when structure is modeled only on the basis of per-nucleotide reactivities (Fig. S8). Moderate DMS reactivities indicate that SL2ext is dynamic, and these dynamics are enhanced in cells, consistent with this region being bound by diverse proteins (Van Herreweghe et al., 2007; Ji et al., 2013; Flynn et al., 2016). Thus, while SL2ext is modeled as lowly probable in cells, the overall consistency between cell-free and in-cell PAIRs leads us to conclude SL2ext is present in state B in cells (Fig. 3D). The interdependence between SL1alt and SL2ext, visualized here, rationalizes prior observations that P-TEFb binding induces structural changes in the 7SK central region, located up to 200 nts away from SL1 (Brogie and Price, 2017). As we discuss below, this structural reorganization overlaps the principal regions bound by P-TEFb release factors, consistent with allosteric coupling between SL1, SL2ext, and release factor binding sites.

Strikingly, RING analysis revealed a dense network of correlations for both cell-free and in-cell RNA (Fig. 3D, S6). Prior *in vitro* studies observed salt-dependent formation of an SL1alt-containing state, consistent with state B potentially being stabilized by tertiary interactions (Brogie and Price, 2017). Some of these RINGs are likely indirect and reflect unresolved minor states. Nonetheless, the consistency and density of observed RINGs suggest that state B contains a compact central core stabilized by dynamic tertiary interactions.

#### State H is a heterogenous P-TEFb released state

The (heterogenous) state H features well-defined SL0, SL1alt, and SL2 stems, but is otherwise highly reactive and contains no other stable structural elements (Fig. 3A, S8). The presence of SL1alt implies that, like B, state H is a P-TEFb released state. However, the elevated reactivities within SL3 and the central domain indicate this state is highly heterogenous. Based on our analysis of simulated data (Fig. S2), we interpret this state to be a composite of diverse lowly populated protein-bound structures. One possibility is that H represents a transition state between A and B that is stabilized by helicases, which are known to bind to regions that are unstructured in H (see Discussion).

#### The 7SK RNA ensemble comprises an allosteric switch

In sum, our data reveal that 7SK folds into at least three structures that comprise P-TEFb– competent and –released states. PAIR and RING measurements, measured individually for each state, provide pivotal and direct evidence of new structural elements (SL2ext) and a compact core in state B that are invisible to per-nucleotide analyses. This multi-state ensemble rationalizes a large compendium of biochemical and functional data on the 7SK RNP, and implies that 7SK contains an allosteric switch that couples HEXIM1/2–P-TEFb binding in SL1 to release factor binding sites in the central domain.

### Mutational analysis validates importance of SL2ext in 7SK structural switching

We validated our DANCE-resolved models and defined the role of individual structural elements in 7SK switching using mutants to probe the A and B states. As an initial control, we performed DANCE-MaP experiments on *in vitro* transcripts of the native sequence RNA. The native RNA folds into a two-state ensemble (Fig. S9), consisting of the A and B states with populations 71 ± 4 and 29 ± 4%, respectively (R > 0.95 between reactivity profiles; N=3). The two states are almost identical to the A and B states observed for the cell-free RNA (Fig. S9A), with the exception that SL0 is further destabilized in State A. The lack of an M state in the *in vitro* transcribed RNA is also consistent with SL0 destabilization. This destablization may reflect increased formal charge at the 5’-phosphate compared the endogenous transcript, which is 5’-*γ*-methylated (Jeronimo et al., 2007; Yang et al., 2019). These data from *in vitro* transcripts further validate that the A and B states are intrinsic features of the 7SK RNA sequence.

To validate the B state structure, we introduced three mismatches in SL1 while leaving SL1alt pairing intact (mutant M1, Fig. 4A,B). DANCE-MaP experiments showed that M1 eliminates state A, with the RNA clustering into 2-3 B or B-like states (N=3; Fig. 4C, S9). Detection of multiple B/B-like states is consistent with the native B state representing a composite of multiple similar structures, and with the M1 mutation modestly destabilizing SL1alt. Rescue of the M1 mutation by restoring base pairing complementarity in SL1 (M1+M2) recovers the native A:B equilibrium (78 ± 2 and 22 ± 2% populations, respectively; N=3; Fig. 4, S9B).

**Figure 4:**
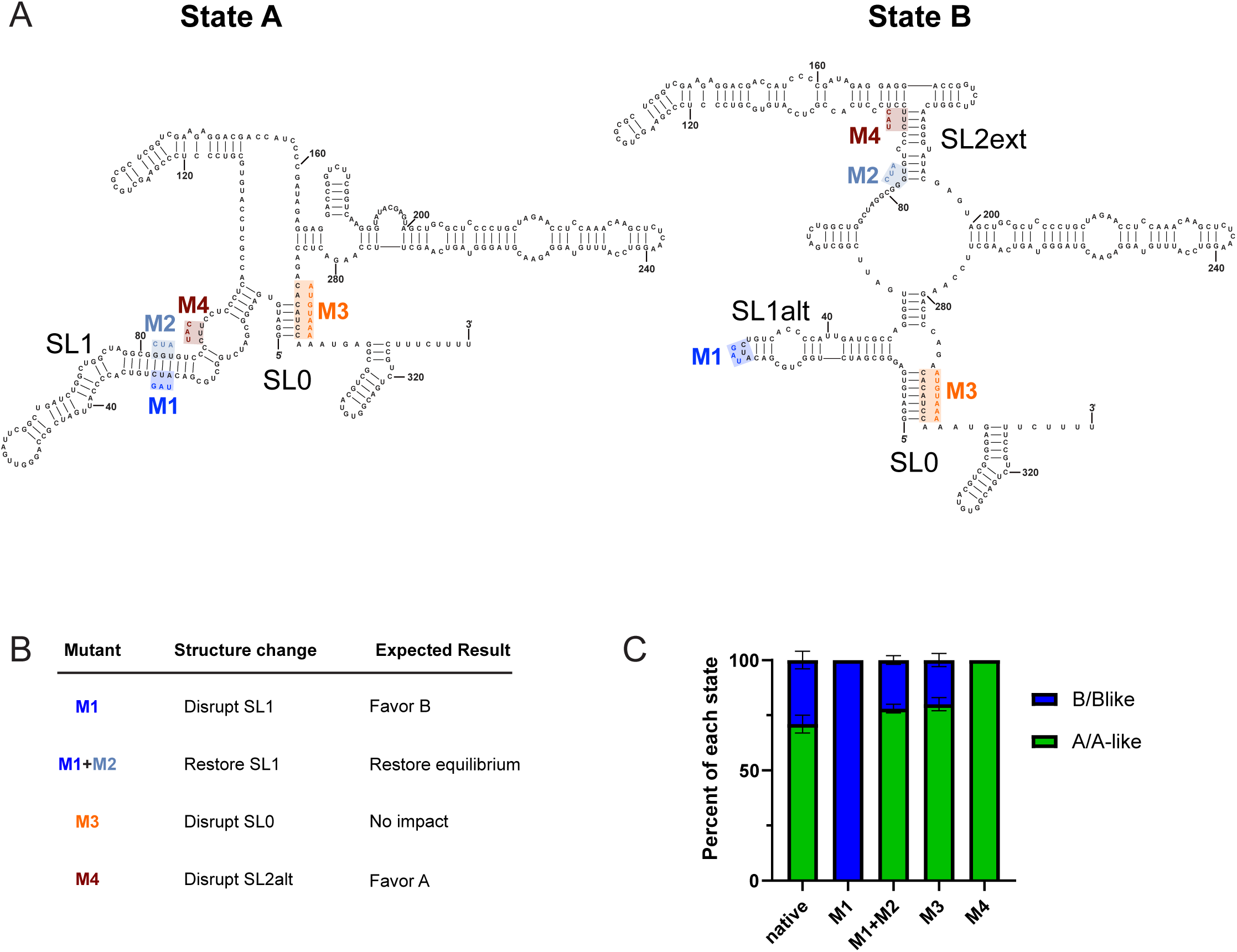
Assessment and validation of 7SK states A and B by mutational analysis. (A) RNA mutants. Mutations are shown superimposed on consensus state A and B structural models. (B) Summary of designed structural impact for each mutant. (C) Ensemble distribution observed for the native sequence RNA and each mutant, produced as *in vitro* transcripts. Structural models are provided in Fig. S9.

We next investigated the role of SL0 in 7SK switching. Our in-cell data support SL0 formation in both states, although SL0 is clearly more stable in state B. Others have proposed that SL0 dynamics drive SL1:SL1alt switching (Brogie and Price, 2017). Ablation of SL0 via the M3 mutation (Fig. 4A) had minimal impact on the 7SK ensemble: M3 populates an 80 ± 3, 20 ± 3% equilibrium consisting of 1-2 A/A-like states and a heterogenous B-like state that replaces SL0 with other interactions (N=3, Fig. S9B). Mutation of only three of seven base pairs in SL0 gave similar results (not shown). Thus, SL0 does not drive the A:B equilibrium.

Finally, we examined the role of SL2ext in A:B switching. PAIR interactions directly support formation of the SL2ext stem for both cell-free and in-cell RNAs (Fig. 3C,D). We thus designed a mutant, M4, to disrupt the PAIR-supported 3-helix junction at the base of SL2ext in state B, but not perturb SL1 pairing in state A (Fig. 4A). Strikingly, this three-nucleotide mutation fully shifts the ensemble to A/A-like states (N=2; Fig. 4C, S9B). Prior studies have also observed that mutations in the SL2ext region induce global remodeling of 7SK structure (Brogie and Price, 2017; Luo et al., 2021), although the mechanistic basis was not resolved. Thus, even though SL2ext shows intermediate stability, this region is critical to 7SK A:B switching.

In sum, the 7SK ensemble can be rationally perturbed via structure-informed mutations, validating the DANCE-resolved A and B states. Moreover, a concise mutation to SL2ext (M4) is sufficient to drive SL1/SL1alt switching, establishing that the central core is an energetically accessible platform for modulating 7SK structure and activity.

### The 7SK structural equilibrium is regulated by cell type and transcriptional demands

The 7SK–P-TEFb axis functions as a global regulator of transcription, and we hypothesized that the 7SK ensemble may be sensitive to cellular gene expression and growth needs. Jurkat cells, like many other tumor cells, exhibit altered P-TEFb regulation and aberrantly upregulated transcription compared to normal cells (Lin et al., 2003; Tyagi et al., 2010). We thus examined whether the 7SK ensemble differs in normal, non-transformed human RPE-1 cells (Bodnar et al., 1998). DANCE-MaP experiments performed on living RPE-1 cells revealed that 7SK adopts precisely the same three structures as observed for Jurkat cells (Fig. 5A) but with significantly different populations: states A, B, and H have populations of 47 ± 2, 34 ± 2, and 19% ± 2, respectively, in RPE-1 cells compared to 40 ± 3, 47 ± 2, 13% ± 1 in Jurkat cells (N=7; Fig. 5B).

**Figure 5:**
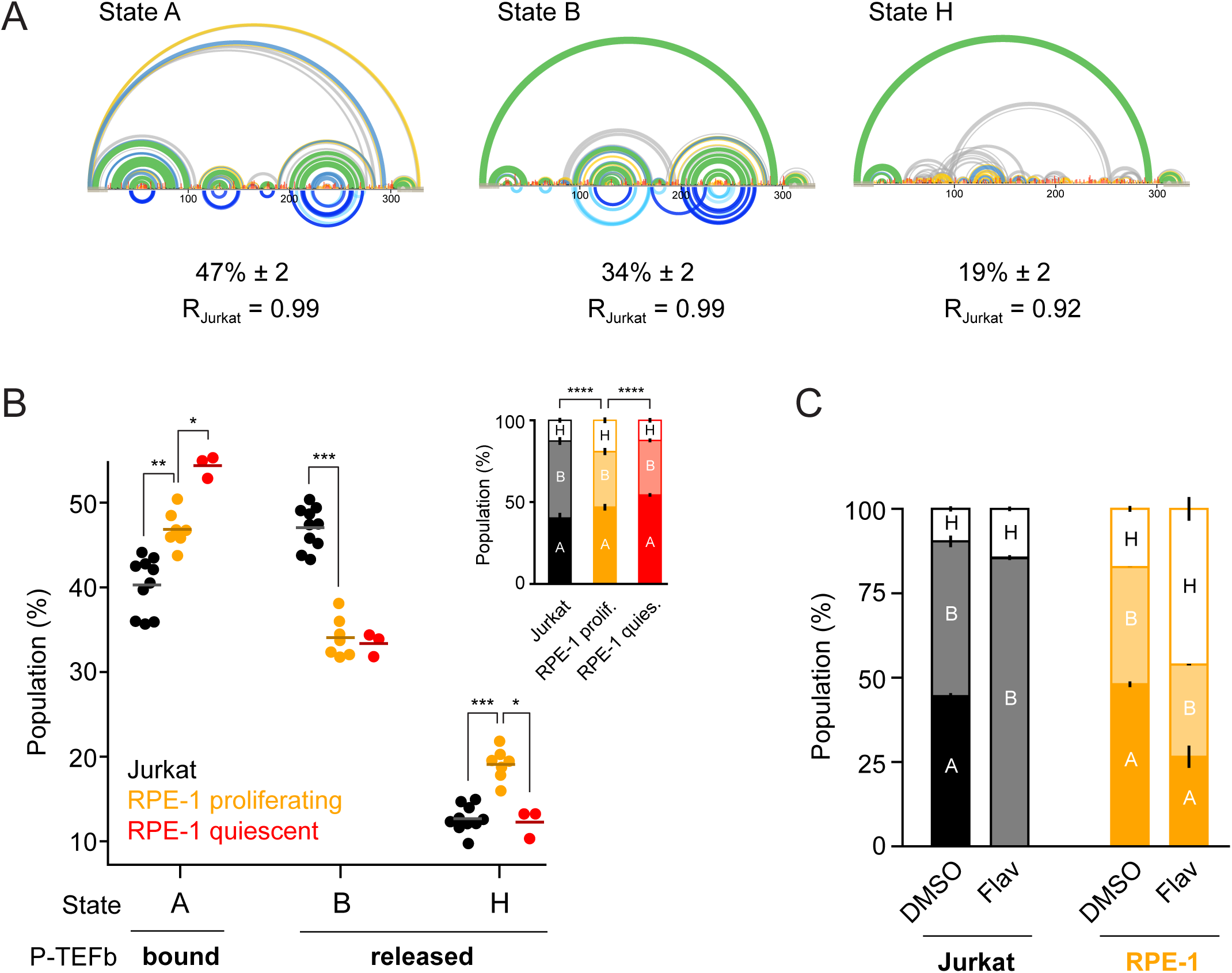
The 7SK equilibrium is regulated by cell type and growth conditions. (A) 7SK structural ensemble in proliferating RPE-1 cells resolved by DANCE-MaP. Structure models are shown as in Figure 3C. Pearson’s R, comparing RPE-1 and Jurkat cell reactivities, are shown. (B) 7SK ensemble populations for Jurkat cells and proliferating and quiescent RPE-1 cells. Comparisons between individual state populations were evaluated using a two-sided Mann-Whitney U test. Inset, population data shown in complete distribution format. Comparisons between complete ensembles were performed using a Dirichlet likelihood ratio test (Shaw et al., 2019). N = 10, 7, and 3 for Jurkat, proliferating RPE-1, and quiescent RPE-1 cells, respectively. *, p<0.05; * *, p<0.01; * * *, p<0.001; * * * *, p<0.0001 (C) Shift in 7SK equilibrium upon flavopiridol treatment in Jurkat and RPE-1 cells. Cells were treated with vehicle (DMSO, 0.01%) or 1 µM flavopiridol for 1 hr (N = 2, all experiments).

This shift represents a ∼30% relative change in the 7SK equilibrium towards A and H and away from B, and is consistent with P-TEFb being more sequestered in RPE-1 cells (or aberrantly released in Jurkat tumor cells). We also probed 7SK in quiescent RPE-1 cells, where transcriptional load is expected to be further decreased. Indeed, the population of state A further increases to 54% ± 1 in contact-inhibited RPE-1 cells (15% relative increase compared to proliferating RPE-1 cells; N=3; Fig. 5B). In absolute molecular terms, each of these changes in 7SK equilibrium correspond to ∼10,000 7SK snRNPs per cell shifting to a P-TEFb binding conformation (Gurney and Eliceiri, 1980; Wassarman and Steitz, 1991).

7SK and P-TEFb are also dynamically regulated in response to transcriptional stress (Peterlin et al., 2012; Quaresma et al., 2016). Flavopiridol is a pan-CDK inhibitor that suppresses transcription by inhibiting CDK9 (Chao et al., 2000). To compensate for reduced CDK9 activity, flavopiridol induces cells to release P-TEFb from 7SK, which conventional probing experiments have indicated induces structural changes in the 7SK RNA (Biglione et al., 2007; Krueger et al., 2010). We directly visualized these structural changes by performing DANCE-MaP experiments in Jurkat and RPE-1 cells treated with either vehicle (DMSO, 0.01%) or 1 µM (saturating) flavopiridol (Biglione et al., 2007). In both cell types, flavopiridol treatment dramatically remodels the 7SK structural ensemble. In Jurkat cells, state A is completely converted to B/B-like states, consistent with total P-TEFb release (Fig. 5C). By contrast, RPE-1 cells exhibit significant, but partial, depopulation of state A (48% to 27%) and conversion to state H (a non-P-TEFb-binding state) (Fig. 5C). This switch supports the physiological relevance of state H and indicates that the 7SK ensemble and P-TEFb release are governed by multiple cell-type-dependent pathways. The attenuated 7SK response to flavopiridol is consistent with the greater tolerance of RPE-1 cells to transcription inhibition as compared to transcription-addicted (Kwiatkowski et al., 2014) Jurkat cells. Collectively, these data establish that the 7SK conformational equilibrium is tunable, is cell type-specific, and remodels dynamically, coincident with P-TEFb release.

### ASO stabilization of state B induces transcription in cells

Our data emphasize that the 7SK structural switch is centrally linked to P-TEFb release and transcription regulation, motivating us to explore the potential of targeting the 7SK ensemble as a strategy for controlling transcription. After screening multiple candidates, we identified an antisense oligonucleotide, ASO-B, that disrupts state A without impacting the major helices unique to state B (Fig. 6A). DANCE-MaP experiments on cell-free RNA confirmed that ASO-B shifted the 7SK structural ensemble to exclusively B/B-like states, whereas a control ASO containing five central mismatches (MM-B) had no significant impact on the 7SK ensemble (Fig. 6B, C). Thus, ASO-B constitutes a molecular tool for modulating the 7SK structural ensemble.

**Figure 6:**
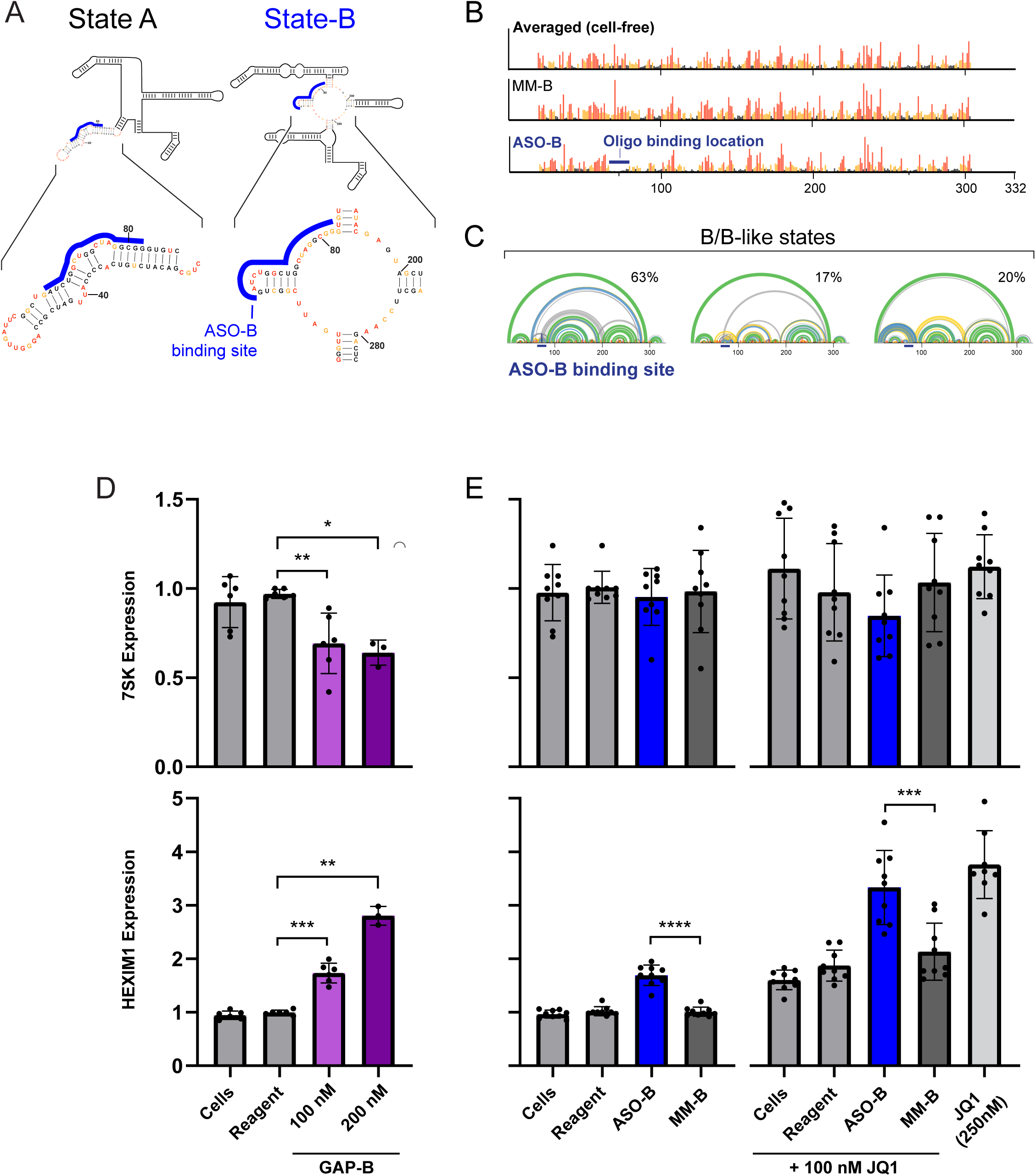
Stabilization of 7SK state B induces transcription. (A) ASO-B binding site shown superimposed on secondary structure models of 7SK states A and B. (B) Engagement of ASO-B on the 7SK RNA observed as a complete reduction of per-nucleotide reactivity at the ASO binding site. MM-B: control ASO containing 5 central mismatches. (C) DANCE deconvolution reveals ASO-B-induced states all correspond to B/B-like states. (D/E) 7SK and HEXIM1 RNA levels measured by RT-qPCR. HEK293 cells were treated for 24 hours with oligos prior to total RNA extraction and quantification. Data normalized to the control gene TBP, in three triplicate experiments. (D) GAP-B induces a decrease in 7SK expression (*top*) and a dose-dependent increase in HEXIM1 expression (*bottom*). N = 3 for 200 nM GAP-B; N = 6 for all other measurements. (E) 7SK expression shows (*top*) no change when treated with ASO-B, MM-B, or JQ1. HEXIM1 shows (*bottom*) significant increase in expression for ASO-B compared to MM-B. N = 9 for all measurements. Significance determined using Welch’s t-test (*, p<0.05; * *, p<0.01; * * *, p<0.001; * * * *, p<0.0001).

We then tested the ability of ASO-B to modulate transcription in cells. We first confirmed ASO delivery and engagement in HEK293 cells using a gapmer oligonucleotide (GAP-B) targeting the same region of 7SK but designed to induce RNase H degradation. Treatment with GAP-B reduced 7SK levels by 31 and 36% at 100 nM and 200 nM concentrations, respectively (Fig. 6D). To assess the impact of 7SK depletion on transcription, we measured the cellular levels of HEXIM1 RNA, which is a sensitive reporter of P-TEFb-mediated transcription activity (He et al., 2006; Liu et al., 2014). As expected, GAP-B-mediated depletion of 7SK induced 1.7-and 2.6-fold induction of HEXIM1 mRNA expression, respectively (Fig. 6D). These data confirm in-cell engagement with 7SK by the ASO-B sequence. We then tested the ability of ASO-B to induce P-TEFb release via stabilization of 7SK state B. Treatment with 100 nM ASO-B but not MM-B yielded a 1.7-fold increase in HEXIM1, without impacting 7SK expression (Fig. 6E), directly validating that 7SK switching induces P-TEFb release. We further examined whether ASO-mediated structure-switching could complement transcriptional activation by the small molecule JQ1. JQ1 is a bromodomain and extra-terminal (BET) inhibitor that induces P-TEFb release via a 7SK-independent mechanism (Chaidos et al., 2014; Fujinaga et al., 2015). Indeed, co-addition of 100 nM JQ1 increased HEXIM1 expression 3.3-fold, comparable to the upregulation observed upon treatment with 250 nM JQ1 alone (Fig. 6E). Together, these experiments further establish a causal relationship between 7SK structural switching and P-TEFb release and provide proof of principle for targeting the 7SK structural switch.

## Discussion

### DANCE-MaP enables complete analysis of RNA structural ensembles

Most RNAs have the potential to fold into multiple structures, which creates numerous opportunities to regulate RNA biology. However, authoritatively defining RNA structural ensembles in cells and their responses to cellular stimuli has remained a critical unresolved challenge. We introduce DANCE-MaP, a single-molecule chemical probing technology that simultaneously measures per-nucleotide reactivities, through-space base pairs (PAIRs), and tertiary interactions (RINGs) for multiple co-existing RNA structural states in cells. DANCE-MaP further measures populations with thermodynamic precision, enabling measurement of ligand binding affinity and of subtle but impactful structural differences between cell types. Collectively, these integrated measurements enable definitive and comprehensive characterization of RNA structural ensembles.

Our study emphasizes the intrinsic complexity of RNA structural ensembles. For the adenine riboswitch, the ON and OFF states are defined by the presence of the aptamer domain and the SD-sequestering helix, respectively, but exhibit substantial heterogeneity elsewhere in the molecule. These data are consistent with and clarify observations from prior biophysical and chemical probing studies (Reining et al., 2013; Warhaut et al., 2017; Tian et al., 2018; Tomezsko et al., 2020). For 7SK, the A, B, and H states are each distinguished by major structural landmarks, but ultimately represent class averages rather than singular conformations. Within this context of underlying heterogeneity, the ability of DANCE-MaP to measure base pairing interactions directly and to estimate pairing probabilities within each state is a crucial advance, enabling us to resolve otherwise invisible dynamic structures in cells and to model global RNA architectures with confidence. Direct PAIR measurements were essential for resolving the SL2ext structure unique to state B of 7SK. RINGs measured in the compact core of state B further emphasize this state has a distinct higher-order structure. Integrated DANCE-MaP analysis represents a powerful advance over single-purpose deconvolution or duplex detection strategies and will broadly enable studies of RNA dynamic complexity in cells.

DANCE-MaP does have several limitations. While our data and other studies (Homan et al., 2014; Mustoe et al., 2019; Tomezsko et al., 2020; Luo et al., 2021; Morandi et al., 2021) support that multiple-hit DMS modification experiments accurately report native RNA structure, accumulated chemical damage may alter behavior of some RNAs. DANCE-MaP can only resolve structural changes that involve >20 nucleotides with populations of at least ∼5%, and currently has a length limit of ∼500 nucleotides across a single strand of RNA. DANCE-MaP has a time-resolution of ∼5 minutes using DMS, but ∼10 second resolution is possible with newer reagents (Ehrhardt and Weeks, 2020). DANCE-MaP further requires that each read originate from a unique RNA molecule, and thus will be more challenging to implement for low abundance RNAs.

Ultimately, DANCE-MaP provides many of the same measurements previously accessible only using state-of-art the NMR experiments, which have provided the primary ground-truth references for RNA ensembles (Liu et al., 2021). The tertiary RINGs measured in the ON state of the adenine riboswitch are of sufficient quality to guide accurate three-dimensional structure modeling (Homan et al., 2014; Li et al., 2020). The RINGs measured in 7SK State B are more challenging to interpret due to the residual dynamics of this state, but such dynamics would similarly challenge established biophysical techniques. Uniquely, DANCE-MaP is readily performed in cells on endogenous RNAs, and requires modest experimental effort. DANCE-MaP thus paves the way for a new generation of biophysical studies in living systems.

### Allostery couples 7SK HEXIM1-P-TEFb aptamer domain to release factor binding sites

Regulated release of P-TEFb from the 7SK snRNP to phosphorylate Pol II is a critical control point in transcription (Peterlin et al., 2012). We show that the 7SK RNA intrinsically encodes a large-scale structural switch that modulates its P-TEFb binding ability (Fig. 7). We further find that the 7SK structural equilibrium is actively controlled by the cell in response to transcriptional demands. Quiescence favors a P-TEFb-sequestered state compared to proliferating cells, and transcriptional stress (CDK9 inhibition) favors a P-TEFb released state (Fig. 5). These population changes correspond to the release or sequestration of tens of thousands of P-TEFb molecules per cell. The amount of free P-TEFb is roughly equivalent to the number of engaged Pol II molecules in a cell (Gurney and Eliceiri, 1980; Kimura et al., 1999; Nguyen et al., 2001; Yang et al., 2001), suggesting that these 7SK/P-TEFb changes play a major role in reshaping global transcription. We also uncover significant changes in the 7SK equilibrium in cancer versus normal cells, providing one of the first examples of cell-type dependence in RNA structure. Collectively, these data support that the 7SK structural switch functions as a key axis of transcriptional regulation.

**Figure 7:**
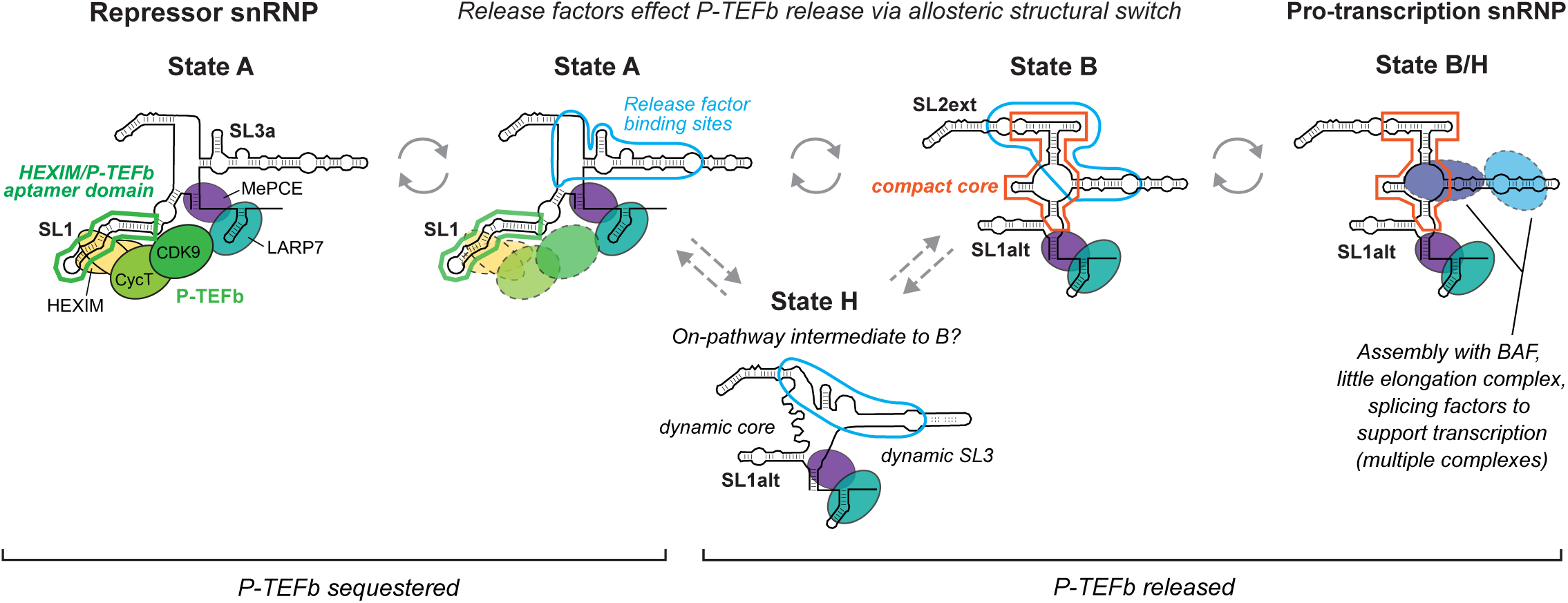
The 7SK ensemble as a dual-function signal integrator. States A, B, H are shown as schematic secondary structures. Compact core corresponds to region of dense RINGs. Annotations show binding sites of core factors (P-TEFb, HEXIM1/2, MePCE and LaRP7), approximate region bound by helicase and other release factors (blue outline), and postulated sites of pro-transcription factors.

Numerous proteins have been implicated as “release factors” that stimulate P-TEFb release from 7SK, a group that includes both helicases and general RNA binding proteins (Van Herreweghe et al., 2007; Ji et al., 2013; Calo et al., 2015; Mück et al., 2016; Bugai et al., 2019; Sithole et al., 2020). Puzzlingly,, these proteins bind 7SK hundreds of nucleotides away from the SL1 hairpin, which comprises the binding site for HEXIM1/2 and P-TEFb (HEXIM/P-TEFb aptamer domain; Fig. 7) (Egloff et al., 2006; Krueger et al., 2008; Martinez-Zapien et al., 2016; Roder et al., 2019). Several studies have observed that HEXIM/P-TEFb release coincides with conformational changes in the SL1 aptamer (Krueger et al., 2010; Brogie and Price, 2017). Binding of hnRNPs has also been observed to induce conformational changes in the 7SK central domain (Luo et al., 2021). However, the relationship between these conformational changes and their role in P-TEFb release remained unclear. Our data resolve this puzzle and support a model whereby release factors catalyze 7SK structural switching and thereby allosterically effect P-TEFb release (Fig. 7). Significantly, release factor binding sites directly overlap or are immediately adjacent to the SL2ext and compact core structures in state B, ideally positioning them to effect structure switching. Given that the unstructured regions of state H also overlap these release factor binding sites, state H may represent an intermediate along the A to B pathway.

This allosteric model of P-TEFb release unifies multiple prior observations, as allostery: (*i*) enables 7SK to maintain a specialized “release domain” that can integrate cellular signals unencumbered by bound P-TEFb; and (*ii*) prevents P-TEFb reassociation once release is triggered. The importance of role (*ii*) is supported by the observation that HIV-1 Tat, despite directly abstracting P-TEFb without binding the 7SK release domain, also induces conformational changes in 7SK (Krueger et al., 2010), which can now be interpreted as switching from state A to B. Less is known regarding how 7SK resequesters P-TEFb. We posit that helicases stimulate disassociation of hnRNPs and remodel 7SK to state A (the SL1-containing form), enabling HEXIM/P-TEFb to rebind. Given that 7SK may be involved in transcription termination (Castelo-Branco et al., 2013), this process may be linked to Pol II recycling. In this model, distinct sets of helicases and other RNA binding proteins catalyze 7SK switching between structural states to either sequester or release P-TEFb.

The allosteric switching model also rationalizes the extreme sequence conservation of the first ∼100 7SK nucleotides across vertebrates and invertebrates (Yazbeck et al., 2018), which must preserve both HEXIM/P-TEFb binding and the dual constraints of forming the distinct SL1 and SL1alt structures. By comparison, the compact core region is highly conserved among Tetrapoda, supporting its functional importance, but diverges outside of Tetrapoda, suggesting that there are multiple ways to create a P-TEFb-regulating allosteric switch. This pattern of a highly conserved P-TEFb aptamer and variable core is also observed in classic riboswitches, where conserved aptamer domains are often integrated with diverse expression domain architectures (Roth and Breaker, 2009).

### 7SK structural switch links P-TEFb release to pro-transcription functions

7SK is canonically considered a transcriptional repressor due to its P-TEFb sequestering function. However, the 7SK snRNP also has pro-transcription functions, including blocking convergent transcription via association with the BAF complex (Flynn et al., 2016) and facilitating spliceosome production (Egloff et al., 2017; Ji et al., 2021). These pro-transcription functions are specific to 7SK snRNPs lacking P-TEFb. Notably, prior chemical probing data obtained for BAF-associated 7SK (Flynn et al., 2016) can now be interpreted as corresponding to state B or H. Together, these data support an overarching model in which the 7SK structural switch integrates P-TEFb release with conversion of 7SK into a pro-transcription snRNP that scaffolds assembly of elongation-supporting factors (Fig. 7, *right*). Significantly, in this model, 7SK switching would enable spatial and temporal coupling between Pol II pause release and BAF-mediated inhibition of convergent transcription (Flynn et al., 2016). Our dual-function model also rationalizes observations that 7SK is inessential for basal P-TEFb regulation (Studniarek et al., 2021), but that 7SK depletion perturbs global chromatin structure (Prasanth et al., 2010) and compromises stress-induced transcriptional reprogramming (Studniarek et al., 2021).

Overall, our model (Fig. 7) emphasizes how structural switching enables the 7SK snRNP to integrate diverse signals to cooperatively inactivate or activate transcription in response to cellular demand. RNAs are unique among biomolecules in their ability to encode large but precise changes in structure (Breaker, 2012; Dethoff et al., 2012). Coupled with the potential of RNAs to form aptamer domains, this makes RNAs optimally suited to serve as molecular integrators. We speculate that similar switching mechanisms broadly underlie non-coding RNA regulatory function.

### 7SK switch constitutes a novel therapeutic target for modulating transcription

Our study shows that the 7SK structural equilibrium is regulated in response to changing transcription needs. We further show using proof-of-principle ASO studies that switching the 7SK state induces transcription of P-TEFb–sensitive targets. Developing small molecules or improved ASOs that stabilize state B, thereby releasing P-TEFb and activating transcription, represents a component of a promising strategy to eradicate persistent HIV infection by inducing the expression of latent provirus (Richman et al., 2009; Cary et al., 2016). Conversely, P-TEFb is commonly dysregulated in cancer and there is intense interest in developing pharmacological inhibitors of P-TEFb as a cancer therapeutic (Yang et al., 2020). Disruption of the 7SK/P-TEFb regulatory axis has been linked to tumorigenesis and cancer progression (Cheng et al., 2012; Ji et al., 2014; Tan et al., 2016), consistent with a model in which dysregulation of the 7SK structural equilibrium supports elevated transcription in cancer cells. Designing small molecules or ASOs that reduce the cellular availability of P-TEFb by selectively stabilizing 7SK state A represents a compelling therapeutic hypothesis for targeting transcription in cancer.

## Acknowledgements

We thank Khoa Dhoa (Baylor College of Medicine, BCM) for exploratory analyses, and Calla Olson (BCM), Kristen Karlin (BCM), and David Price (University of Iowa) for helpful discussions. Research reported in this publication was supported by Qura Therapeutics (D.M.M.), by the NIH (R35 GM122532 to K.M.W), the Cancer Prevention & Research Institute of Texas (RR190054 to A.M.M.), and BCM seed funds (to A.M.M.). The content of this study is solely the responsibility of the authors and does not necessarily represent the official views of Qura Therapeutics.

## Disclaimer

A.M.M. is a consultant to, and K.M.W. is an advisor to and holds equity in Ribometrix.

