## Supporting Information for "Large-scale allosteric switch in the 7SK RNA regulates transcription in response to growth and stress"

Methods, Supporting Figures (10), and Supporting Table (1)

#### Methods

##### DANCE-MaP algorithm

Data pre-processing. Aligned DMS-MaP sequencing reads are processed into two binary vectors. The mutation string, $\boldsymbol{x_n} = (x_{n,1}, \dots, x_{n,D})$, encodes whether a match (0) or a mutation (1) is observed at each nucleotide, where *n* indexes an individual read and *D* is the length of the amplicon. The data string, $\boldsymbol{\delta_n} = (\delta_{n,1}, \dots, \delta_{n,D})$, encodes whether the nucleotide was measured (1) or not (0); example "no measurement" scenarios include low quality score, a deletion, or masking due to minimum spacing requirements between mutations (Busan and Weeks, 2018). Reads with fewer than 75% of positions defined, $\sum_i^D \delta_{n,i} < 0.75 \cdot D$, are discarded. Positions with mutation rates >0.02 in the ethanol-treated control sample ($r_{etoh,i}$), or with average mutation rates less than 0.0001 in the DMS-treated sample ($r_{DMS,i}$), are considered invalid and excluded from further analysis ($\delta_{n,i} = 0$ for all *n*).

To prevent individual nucleotides from dominating the maximum likelihood (ML) clustering outcome and to improve clustering power, nucleotide positions are split into "active" and "inactive" categories. Active nucleotides are included in likelihood calculations during primary ML clustering, whereas inactive nucleotides are excluded and solved via a second constrained ML-optimization (Fig. S1). Active versus inactive status is specified using the binary vector $\boldsymbol{\phi} = (\phi_1, \dots, \phi_D)$, where $\phi_i = 1$ denotes active status. Nucleotides where $(r_{DMS,i} - r_{etoh,i})$ < 0.002 are set to inactive upon model initialization. As described below, additional nucleotides may be inactivated over the course of model solution.

Primary ML clustering. Data are fit to a Bernoulli mixture model using the expectation-maximization (EM) algorithm (Bishop, 2006). Fitting is performed for sequentially larger numbers of model components (structural states), beginning with 1, until the best fit is identified (Fig. S1).

A model consisting of K components is specified by two parameter vectors:

the populations of each state, $\boldsymbol{\pi} = (\pi_1, \dots \pi_K)$ ; $\sum \pi_k = 1$

and the mutation rates of each state, $\boldsymbol{\mu} = \begin{pmatrix} \mu_{1,1}, \dots \mu_{1,D} \\ \vdots \\ \mu_{K,1}, \dots \mu_{K,D} \end{pmatrix}$

The fitting process begins with random initialization of $\boldsymbol{\mu}$ (drawn from {Beta(1,40)+0.001}) and $\boldsymbol{\pi} = (1/K, \ldots\ 1/K)$. Solutions for $\{\boldsymbol{\pi}, \boldsymbol{\mu}\}$ are then obtained via EM iteration. The appropriate equations for EM iteration incorporating missing data and active/inactive positions are:

$$z_{n,k} = \frac{\pi_k \prod_{i=1}^{D} \left[\mu_{k,i}^{x_{n,i}} (1-\mu_{k,i})^{1-x_{n,i}}\right]^{\phi_i\, \delta_{n,i}}}{\sum_{m=1}^{K} \pi_m \prod_{i=1}^{D} \left[\mu_{m,i}^{x_{n,i}} (1-\mu_{m,i})^{1-x_{n,i}}\right]^{\phi_i\, \delta_{n,i}}} \qquad [1]$$

$$\mu_{k,i} = \frac{\sum_{n=1}^{N} z_{n,k}\, \delta_{n,i}\, x_{n,i} + \alpha_i - 1}{\sum_{n=1}^{N} z_{n,k}\, \delta_{n,i} + \alpha_i + \beta_i - 2} \quad for\ \ i \in \{active\} \qquad [2.1]$$

$$\pi_k = \frac{\sum_{n=1}^{N} z_{n,k}}{N} \qquad [2.2]$$

*N* is the total number of reads being clustered. $\alpha_i$ and $\beta_i$ are parameters for a nucleotide-specific beta prior, which are set to:

$$\alpha_i = 1 + 0.01 \cdot r_{etoh,i} \cdot \textstyle\sum_{n=1}^{N} \delta_{n,i}\ ;\ \ \beta_i = 2$$

This prior encourages convergence to solutions with $\mu_{k,i} \geq r_{etoh,i}$ (preventing convergence to non-physical solutions where $\mu_{k,i} \to 0$), and contributes approximately $^1/_{100}$ the weight relative to the input data in determining the final value of $\mu_{k,i}$.

Equations [1] and [2.1, 2.2] are iterated until convergence, defined as $\max\left(\left|\boldsymbol{\mu}^{(t+1)} - \boldsymbol{\mu}^{(t)}\right|\right) < 10^{-4}$ and $\max\left(\left|\boldsymbol{\pi}^{(t+1)} - \boldsymbol{\pi}^{(t)}\right|\right) < 10^{-4}$, where $\{\boldsymbol{\pi}^{(t+1)}, \boldsymbol{\mu}^{(t+1)}\}$ denotes the parameters at iteration *t+1.* Converged solutions are assessed for validity as defined below. If an invalid solution is obtained repeatedly, then the nucleotides causing the invalid failure are inactivated ($\phi_i = 0$).

EM fitting is repeated from different random initializations until 3 identical converged solutions are found, up to a maximum of 50 attempts. Identical solutions are defined as $\max\left(\left|\boldsymbol{\pi}^a - \boldsymbol{\pi}^b\right|\right) < 0.03$ and $\max\left(\left|\boldsymbol{\mu}^a - \boldsymbol{\mu}^b\right|\right) < 0.01$, where *a* and *b* denote two different valid K-component solutions. If 3 solutions are not identified, then the model search terminates, selecting the K-1 model. If 3 identical solutions are identified, then the Bayesian information criteria (BIC) is used to evaluate whether the K-component model is significantly better than the K-1 model. Specifically, we compute:

$$\Delta BIC = (q_K - q_{K-1})\ln(N) - 2\ln\left(\mathcal{L}_K / \mathcal{L}_{K-1}\right)$$

where $q_K$ and $\mathcal{L}_K$ denote the number of parameters and the likelihood of the *K*-component model, respectively. If $\Delta BIC \leq -46$, indicating ~$10^{10}$ greater evidence for the *K* versus *K-1* model, then the *K*-component model is accepted, *K* is incremented, and the EM fitting process repeats to a default maximum of *K*=5. Otherwise, the *K-1* model is selected and the search terminates. We denote the final selected parameters as $\{\hat{\boldsymbol{\pi}}, \hat{\boldsymbol{\mu}}\}$.

Note that final fitted parameters can occasionally exhibit small run-to-run variation, caused by different sets of nucleotides being inactivated during each stochastic fitting trajectory. When observed, this variation is within the expected precision of the overall experiment: $\hat{\boldsymbol{\pi}}$ correlate with R>0.995, and $\hat{\boldsymbol{\mu}} \pm 3\%$.

Model validity criteria. Initial testing revealed that the EM algorithm occasionally converged to non-physical or degenerate solutions, particularly for K>2. We thus we perform several tests to ensure model validity; solutions that fail any of these validity criteria are rejected.

First, we require that $\{\boldsymbol{\pi}, \boldsymbol{\mu}\}$ fall within the physical boundaries:

$$\min(\boldsymbol{\pi}) \geq 0.001$$
$$\min(\boldsymbol{\mu}) \geq 10^{-5}$$
$$\max(\boldsymbol{\mu}) \leq \min(0.5, \quad 3 \cdot \max(\boldsymbol{r_{DMS}}))$$
$$\mu_{k,i} / \mu_{l,i} \leq 200 \text{ for } i \in \{active\};\ k,l \in \{1, \dots K\},\ k \neq l$$

Second, to prevent selection of degenerate models, we require the root mean square (RMS) difference between reactivity profiles be ≥ 0.005, excluding the top percentile of $\boldsymbol{\mu}$ differences:

$$RMS_{k,l} = \sqrt{\langle(\mu_{l,i} - \mu_{k,i})^2\rangle_{i\in\{active\};\,(\mu_{l,i}-\mu_{k,i})^2 \leq P_{99}}} \geq 0.005 \text{ for all } k,l \in \{1, \dots K\},\ k \neq l$$

where $P_{99}$ denotes the 99th percentile of $(\mu_{l,i} - \mu_{k,i})^2$ values.

Third, because the minimum spacing required between mutations can favor artifactual solutions featuring anticorrelated $\boldsymbol{\mu}$ parameters over short-length scales, we discriminate against such solutions by computing the scores

$$s_{1,i} = \sum_{m=0}^{2} \phi_{i+m} \log\left(\frac{\mu_{k,i+m}}{\mu_{l,i+m}}\right) + \sum_{m=3}^{7} \phi_{i+m} \log\left(\frac{1-\mu_{k,i+m}}{1-\mu_{l,i+m}}\right) + \sum_{m=8}^{10} \phi_{i+m} \log\left(\frac{\mu_{k,i+m}}{\mu_{l,i+m}}\right)$$

$$s_{2,i} = \sum_{m=0}^{2} \phi_{i+m} \log\left(\frac{1-\mu_{l,i+m}}{1-\mu_{k,i+m}}\right) + \sum_{m=3}^{7} \phi_{i+m} \log\left(\frac{\mu_{l,i+m}}{\mu_{k,i+m}}\right) + \sum_{m=8}^{10} \phi_{i+m} \log\left(\frac{1-\mu_{k,i+m}}{1-\mu_{l,i+m}}\right)$$

for all positions *i*. Solutions are rejected if $|s_{1,i}| > 4.6$ and $|s_{2,i}| > 4.6$ for any *i*, equivalent to a >100-fold anticorrelated likelihood difference over a 10-nt window.

Finally, to prevent selection of poorly defined models, we require that the information matrix for $\{\boldsymbol{\pi}, \boldsymbol{\mu}\}$ be invertible. The information matrix is computed from the observed data as described (McLachlan and Peel, 2000).

Solving for inactive positions. Following identification of a valid converged solution, $\mu_{k,i}$ parameters of inactive nucleotides are obtained via constrained EM fitting. $\pi_k$ and active $\mu_{k,i}$ are held fixed while iterating the equations

$$z'_{n,k} = \frac{\pi_k \prod_{i=1}^{D} \left[\mu_{k,i}^{x_{n,i}} (1-\mu_{k,i})^{1-x_{n,i}}\right]^{\delta_{n,i}}}{\sum_{m=1}^{K} \pi_m \prod_{i=1}^{D} \left[\mu_{m,i}^{x_{n,i}} (1-\mu_{m,i})^{1-x_{n,i}}\right]^{\delta_{n,i}}}$$

$$\mu_{k,i} = \frac{\sum_{n=1}^{N} z'_{n,k}\, \delta_{n,i}\, x_{n,i} + \alpha_i - 1}{\sum_{n=1}^{N} z'_{n,k}\, \delta_{n,i} + \alpha_i + \beta_i - 2} \quad for\ \ i \in \{inactive\}$$

until the inactive $\mu_{k,i}$ converge (defined as $\max\left(\left|\boldsymbol{\mu}^{(t+1)} - \boldsymbol{\mu}^{(t)}\right|\right) < 10^{-4}$ ).

Final model quality assessment. *In silico* benchmarking revealed that ML fitting occasionally yielded “valid” solutions that were nonetheless inaccurate in relation to the known generating ensemble. These inaccurate solutions were reliably identified as having poorly differentiated reactivity profiles, with relatively few nucleotides differing between model states. Thus, we perform a final quality assessment and warn users of potential low-quality solutions. These quality checks consist of:

i) Confirming that the RMS difference between all profiles is sufficiently different,

$$RMS_{k,l} = \sqrt{\langle(\hat{\mu}_{l,i} - \hat{\mu}_{k,i})^2\rangle_{i\,\in\,active}} \;\; \geq 0.01 \text{ for all } k, l \in \{1, \ldots K\},\ \ k \neq l$$

ii) Confirming that at least 20 nucleotide positions in each profile have distinct reactivities by computing the number distinct score (ND)

$ND_{k,l} = \left[\sum_i \phi_i \cdot \mathbf{1}_A\left(\left|\hat{\mu}_{l,i} - \hat{\mu}_{k,i}\right| > 0.01\right)\right] \geq 20$ for all $k, l \in \{1, \ldots K\},\ k \neq l$

where $1_A$ is the indicator function.

iii) Confirming that the populations are well-defined according to the information matrix

$$\max\left(\sigma(\hat{\pi}_k)\right) < 0.01$$

where $\sigma(\hat{\pi}_k)$ is the standard deviation of $\hat{\pi}_k$ obtained from the square root of the inverse of the information matrix.

Reactivity normalization. Following clustering, $\hat{\boldsymbol{\mu}}$ parameters are transformed into normalized DMS reactivities that can be used as input for structure modeling. Raw reactivity profiles ($r_{raw,k}$) for each state *k* are obtained by subtracting $r_{etoh}$ from $\hat{\boldsymbol{\mu}}$. Nucleotide-specific normalization factors ($n_i$) that scale A/C and U/G reactivities to similar 0 to ≈2 ranges are computed as described (Mustoe et al., 2019) based on the maximum raw reactivities observed over all states. Normalized reactivities ($r_k$) are then obtained as $r_{raw,k}/n_i$.

RING- and PAIR-MaP analysis. Given a converged Bernoulli mixture model $\{\hat{\boldsymbol{\pi}}, \hat{\boldsymbol{\mu}}\}$, individual reads can be assigned *a posteriori* to the component (structure) from which they were derived. These assigned reads can then be input to PAIR and RING analyses, which identify correlated modifications between pairs of nucleotides that are indicative of through-space base pairing and tertiary interactions (Homan et al., 2014; Mustoe et al., 2019).

In principle, each read *n* can be assigned to its most probable parent structure using the complete data vector [$\mathbf{x_n} = (x_{n,1}, x_{n,2}, \ldots, x_{n,D})$]. However, in practice, the modification status of a nucleotide ($x_{n,i}$) can strongly bias read assignment and thereby impose correlations in the assigned data. We address this issue by performing read assignment independently for each pair of nucleotides, excluding the considered nucleotides from the posterior probability calculation (Fig. S1). Specifically, for the nucleotide pair (*v*, *w*), the posterior probability of a read $\mathbf{x_n}$ being derived from a structure *k* is computed as

$$z_{n,k}(v,w) = \frac{\hat{\pi}_k \prod_{\forall\, i \neq v,w} \left[\hat{\mu}_{k,i}^{x_{n,i}}(1-\hat{\mu}_{k,i})^{1-x_{n,i}}\right]^{\delta_{n,i}}}{\sum_{m=1}^{K} \hat{\pi}_m \prod_{\forall\, i \neq v,w} \left[\hat{\mu}_{m,i}^{x_{n,i}}(1-\hat{\mu}_{m,i})^{1-x_{n,i}}\right]^{\delta_{n,i}}}$$

Only reads that can be confidently assigned to a parent structure, defined as $\{\mathbf{X_k}(v,w)\} = \{\mathbf{x_n} \mid z_{n,k}(v,w) \geq 0.9\}$, are used for correlation analysis. The modification status of *v* and *w* is then tabulated across $\{\mathbf{X_k}(v,w)\}$ to obtain the {(*unmod*, *unmod*), (*mod*, *unmod*), (*unmod*, *mod*), (*mod*, *mod*)} contingency table (Homan et al., 2014; Mustoe et al., 2019). Note that this scheme means that the same read $\mathbf{x_n}$ can be assigned to different parent structures for different (*v, w*). For PAIR-MaP analysis, which considers correlations between windows of 3 nucleotides, nucleotides (*v, v+1,* v+2*, w, w+1, w+2*) are excluded from the products in the $z_{n,k}(v,w)$ equation.

As an additional control to eliminate artifactual correlations from biased read assignment, we perform an identical read-assignment and correlation analysis on a matched uncorrelated synthetic dataset (Fig. S1). The synthetic dataset is generated from the experimentally fitted $\hat{\pi}_k$ and $\hat{\mu}_{k,i}$ parameters, treating all nucleotides as independent, and contains an identical number of reads as the experimental dataset. Any nucleotide pairs (*v, w*) observed correlated in this null dataset (P<0.001, G-test) are removed from the set of experimental correlations. Additionally, the contingency table for each (*v, w*) pair measured for the experimental data is required to be significantly different than the contingency table obtained for the null dataset (P<0.001, G-test); (*v, w*) pairs that fail this test are likewise removed from the set of experimental correlations.

In developing the read-assignment algorithm, we also tested the following alternative strategies: assigning reads using the complete data vector (not excluding (*v, w*)); using maximum *a posteriori* assignment; and assigning reads using Monte Carlo selection. Benchmarking tests on synthetic datasets unequivocally showed the strategy described above to have the best sensitivity and specificity.

After read assignment, RING and PAIR analysis are performed using v1.1 of RingMapper (Mustoe et al., 2019).

**Cell culture**

Jurkat E6-1 cells were obtained from ATCC (TIB-152) and cultured in suspension using RMPI 1640 media (Gibco) supplemented with 10% FBS (Millipore), 100 U/mL Pen/Strep (LifeTech) at 37 °C and 5% $CO_2$. hTERT RPE-1 (RPE-1) cells were a gift from W. Marzluff (UNC) and were authenticated by STR profiling and confirmed to be free of mycoplasma contamination. RPE-1 cells were maintained in DMEM/F-12 + HEPES (Gibco) with 10% FBS (Gibco), 100 U/mL Pen/Strep (Gibco), 2 mM sodium pyruvate (Gibco), and MEM non-essential amino acids (Gibco) at 37 °C and 5% $CO_2$. HEK293T/17 cells were obtained from ATCC (CRL-11268) and maintained in DMEM (LifeTech) supplemented with 10% FBS (Millipore) and 100 U/mL

Pen/Strep at 37 °C and 5% $CO_2$.

**DMS probing of the adenine riboswitch RNA**

Native sequence and mutant *V. vulnificus add* adenine riboswitches containing 5' and 3' structure cassettes were transcribed *in vitro* (Mustoe et al., 2019). Briefly, templates were synthesized as gBlocks [IDT; (Mustoe et al., 2019) and Table S1], amplified by PCR (Q5 DNA polymerase, NEB), and purified (PureLink PCR column, Invitrogen). RNA was transcribed *in vitro* [400 µL; 40 mM Tris (pH 8.0), 25 mM $MgCl_2$, 2.5 mM Spermidine, 0.01% (vol/vol) Triton X-100, 10 mM DTT, 5 mM each NTP, ~4 µg DNA template, 0.05 mg/mL T7 RNA polymerase (lab made), 0.2 U pyrophosphatase (NEB); 37 °C; 4h], treated with DNase (TURBO DNase, Invitrogen), purified (Agencourt RNAclean XP beads; Beckman Coulter), and stored at -20 °C. RNA size and purity was confirmed using Bioanalyzer analysis (Agilent) and concentration was quantified (Qubit RNA BR assay, Invitrogen).

For probing experiments, RNA [4 pmol in 2 µL volume] was denatured at 95 °C for 2 min followed by snap cooling on ice for 2 min. RNA was folded by adding 7 uL of 1.43× adenine-containing folding buffer [1× buffer: 300 mM bicine (pH 8.0), 100 mM NaCl, 5 mM $MgCl_2$, variable adenine] and incubated at 30 °C for 30 min. Folded RNA was added to 1 µL of DMS solution (1.7 M in ethanol), allowed to react for 10 min at 30 °C, and then quenched via addition of an equal volume of 20% 2-mercaptoethanol (vol/vol in $H_2O$) and placed on ice. RNA was purified by precipitation with ethanol. No-reagent control RNA was prepared identically, substituting neat ethanol for the DMS solution.

**DMS probing of 7SK RNA in cells**

For Jurkat cells, cells were pelleted, washed with PBS, and counted. 1-2×$10^6$ cells were resuspended in 450 µL fresh media supplemented with 200 mM Bicine (pH 8.0). Cells were then treated with 50 µL of 1.7 M DMS in ethanol or 50 µL ethanol for 6 min at 37 °C. Reactions were quenched with 500 µL 20% 2-mercaptoethanol and placed on ice. Cells were pelleted and RNA extracted using 1 mL TRIzol reagent (Invitrogen). Residual DNA was removed by treating with 2 units of TURBO DNase (Ambion) for 30 min at 37 °C, followed by spike-in of 2 additional units and further 30 min incubation (1 hour total). RNA was purified by SPRI beads (MagBind TotalPure NGS beads; Omega BioTek) and quantified by UV absorbance (Nanodrop).

For RPE-1 cells, 1.5×$10^6$ cells were seeded into a 10 cm dish 48 hr prior to probing. Media was removed and 5.4 mL fresh media, supplemented with 200 mM Bicine (pH 8.0), was added and incubated at 37 °C for 3 min. Cells were treated with 600 µL of 1.7 M DMS or neat ethanol for 6 min at 37 °C, followed by quenching using 6 mL of 20% 2-mercaptoethanol on ice. Cells were scraped and pelleted, RNA was extracted using TRIzol (as described for Jurkat cells) or column (RNeasy mini; Qiagen), and quantified by UV absorbance (Nanodrop).

**DMS probing of cell-free 7SK RNA**

Total RNA was extracted from $2\times10^6$ Jurkat cells using TRIzol reagent (Invitrogen). RNA was DNase treated, purified (Mag-Bind TotalPure NGS beads; 1.8× ratio), and quantified as described above for in-cell RNA. 2 µg RNA in 50 µL in water was denatured at 98 °C for 1 min, snap cooled at 4 °C for 1 min, and then refolded via addition of 50 µL of 2× Bicine RNA folding buffer and incubation at 37 °C for 20 minutes [1× folding buffer: 200 mM Bicine (pH 8.0), 200 mM potassium acetate (pH 8.0) and 5 mM $MgCl_2$] (Mustoe et al., 2019). Samples were split into two 45 µL aliquots and treated with either 5 µL 1.7 M DMS in ethanol or 5 µL neat ethanol at 37 °C for 6 minutes. Following treatment, samples were quenched with 1 volume of 20% 2-mercaptoethanol, placed on ice, and purified by isopropanol precipitation.

**DMS probing of *in vitro* transcribed 7SK RNA**

DNA templates were synthesized as gBlocks (Integrated DNA technologies; Table S1) and amplified by PCR [Q5 HotStart polymerase (NEB), supplemented with 1.0 M betaine]. DNA templates were purified (Mag-Bind TotalPure NGS beads; 0.7× ratio). RNA was transcribed *in vitro* [400 µL; 40 mM Tris (pH 8.0), 25 mM $MgCl_2$, 2.5 mM spermidine, 0.01% (vol/vol) Triton X-100, 10 mM DTT, 5 mM each NTP, 200 ng DNA template, 95 µg T7 RNA polymerase (lab made), 20 U RNasin (Promega), 50 U yeast inorganic pyrophosphatase (NEB); 37 °C; 4h]. Transcription reactions were treated with 16 U TURBO DNase (Thermo) for 30 min at 37 °C and purified (Mag-Bind TotalPure NGS beads; 1.8× bead:volume ratio) and stored at -20 °C. RNA size and purity were confirmed using Bioanalyzer analysis (Agilent) and concentration was quantified (Qubit RNA BR assay, Invitrogen).

For probing experiments, RNA [10 µg in 50 µL] was denatured at 95 °C for 2 min followed by snap cooling on ice for 2 min. 50 uL of 2× folding buffer was then added and the RNA folded at 37 °C for 30 min [1× buffer: 200 mM Bicine (pH 8.0), 200 mM potassium acetate (pH 8.0) and 5 mM $MgCl_2$]. 45 µL of folded RNA was added to 5 µL of DMS solution (1.7 M in ethanol), allowed to react for 6 min at 37 °C, quenched via addition of an equal volume of 20% 2-mercaptoethanol, and placed on ice. RNA was purified by precipitation with isopropanol. No-reagent control RNA was prepared identically, substituting neat ethanol for the DMS solution.

**DMS probing of contact-inhibited cells**

RPE-1 cells were growth-arrested by contact inhibition as described (Matson et al., 2019). For each replicate experiment, three 10 cm dishes were seeded with $6 \times 10^6$ RPE-1 cells and grown to 100% confluency. Medium was then exchanged and the cells were incubated for an additional 96 hr to allow for complete growth arrest. Two dishes of arrested cells were modified with DMS or ethanol as described for proliferating RPE-1 cells. The third dish was used to confirm growth arrest by flow cytometry analysis. Cells were incubated with 10 µM 5-ethynyl-2´-deoxyuridine (EdU) for 1 hr prior to harvesting. Cells were then harvested with trypsin and fixed in 4% formaldehyde. Cells were labeled with Alexa Fluor 488–azide and nuclei were stained with DAPI. Less than 1% of cells were in S phase (positive for Alexa 488), and 90% of cells were in G1/G0 (2*n* DNA content measured by DAPI).

**DMS probing of flavipiridol treated cells**

Jurkat cells (3 million cells in 10 mL fresh media) were seeded 23 hours prior to treatment and were then treated with either vehicle (0.01% DMSO) or with 1 µM flavopiridol (in DMSO) for 1 hour. RPE-1 cells were seeded 23 hr prior to be 70% confluent on the day of experiment and were treated with 0.01% DMSO or 1 µM flavopiridol for 1 hour. Cells were then treated with DMS and RNA was extracted identically as described above for in-cell probing experiments.

**MaP reverse transcription**

Mutational profiling (MaP) reverse transcription (RT) was performed exactly as described (Mustoe et al., 2019; Sengupta et al., 2019). For adenine riboswitch experiments, one-half of the purified DMS reaction was input into RT. For in-cell and cell-free 7SK experiments, 1 µg total cellular RNA was input into RT. For *in vitro* 7SK experiments, 100 ng RNA was input into RT. RT products were purified by beads (Mag-Bind TotalPure NGS beads; 1.8× ratio) or column (G-50 Sephadex; Cytiva).

**Sequencing library construction**

Sequencing libraries were generated using the two-step PCR approach (Smola et al., 2015a).

For the adenine riboswitch, one-seventeenth of the purified RT reaction was input to PCR1, performed [98 °C for 30 s, 10 cycles of (98 °C for 8 s, 66 °C for 20 s, 72 °C for 20 s), and 72 °C for 2 min]. PCR1 product was purified (Mag-Bind TotalPure NGS beads; 0.8× ratio). 2.5 ng product was input to PCR2 [98 °C for 30 s, 10 cycles of (98 °C for 8 s, 68 °C for 20 s, 72 °C for 20 s), and 72 °C for 2 min]. PCR2 product was purified (Mag-Bind TotalPure NGS beads; 0.8× ratio), and sequenced with an Illumina MiSeq instrument using 2×150 paired-end sequencing (v2 chemistry). Data used for high-depth PAIR-MaP/RING-MaP analysis in Figure 2 were obtained by resequencing libraries from a previously published experiment (Mustoe et al., 2019).

For 7SK, one-fifth of the purified RT reaction was input to PCR1 [98 °C for 30 s, 10 cycles of (98 °C for 10 s, 68 °C for 20 s, 72 °C for 20 s), and 72 °C for 2 min]. PCR1 product was purified (Mag-Bind TotalPure NGS beads; 0.8× ratio). 1-2 ng product was input to PCR2 [98 °C for 30 s, 10-14 cycles of (98 °C for 10 s, 65 °C for 30 s, 72 °C for 20 s), and 72 °C for 2 min]. PCR2 product was purified (Mag-Bind TotalPure NGS beads; 0.8× ratio) and sequenced with an Illumina MiSeq instrument using 2×250 (v2 chemistry) or 2×300 (v3 chemistry) paired-end sequencing.

**Sequence alignment and data analysis**

*ShapeMapper* (v2.1.5) was used to align and parse mutations from DMS-MaP sequencing experiments using the --amplicon and --output-parsed-mutations options. Adenine riboswitch data were aligned against the synthesized template sequence, and 7SK data were aligned against NR_001445.2. DANCE-MaP analysis was performed using the *DanceMapper* (v1.0) software. For the adenine riboswitch, *DanceMapper* was run with default options, which allows clustering into a maximum of 5 components. For 7SK, *DanceMapper* was run allowing a maximum of 3 clusters (--maxc=3); the absence of higher-order clustering solutions was confirmed via analysis of selected datasets. PAIR and RING analyses were performed via *DanceMapper* using default options.

**7SK replicate analyses**

7SK per-nucleotide reactivity data, PAIRs, RINGs, and structure models are derived from single experiments and are representative of at least 2 independent replicate datasets and analyses. Note that the sensitivity of PAIR and RING analysis depends strongly on read depth. The high-depth Jurkat in-cell and cell-free datasets shown in Figure 3/S6 were deliberately sequenced to high depths (>3 million), and the comparative lack of PAIRs and RINGs in other samples is attributable to lower sequencing coverage (0.3 to 1 million reads per sample). Ensemble populations are reported as the mean and standard deviation across replicates.

For cell-free Jurkat probing experiments, a total of 4 independent replicates were collected. Data did not reliably cluster into 3 states at read-depths below 1 million (2 of 4 replicates clustered into two-state ensembles consisting of A and B). Population means and errors were thus computed by comparing results from the single deeply sequenced dataset to a consolidated replicate constructed from the remaining 3 replicates. This consolidated replicated was also used to assess PAIR and RING reproducibility in Figure S6.

PAIR and RING reproducibility for in-cell Jurkat data were assessed by pooling reads from 8 independent replicates to create the "consolidated replicate" shown in Figure S6.

**Structure modeling**

Structure modeling was performed using *RNAstructure* (v6.2) (Reuter and Mathews, 2010). The *partition* module was modified to enable DMS-guided pairing probably calculations using nucleotide-specific DMS reactivity restraint functions (Mustoe et al., 2019); this modified code is available upon request and will be distributed in future releases of *RNAstructure*. Normalized DMS and PAIR restraints output by *DanceMapper* were passed to *fold* and *partition* using the –dmsnt and –x flags, respectively. Pairing probabilities shown in Figure 1C were computed using DMS reactivities only. All other structure modeling was performed using both DMS reactivities and PAIR restraints (when available). As part of *DanceMapper*, we provide the script *foldClusters*.py that automates structure modeling and visualization for all states of a deconvoluted ensemble.

**ASO experiments**

The ASO-B antisense oligonucleotide was designed to bind 7SK nts 64-82 to stabilize state B and contained complete 2'-O-methyl modifications to render it insensitive to RNase H. The mismatch MM-B ASO contains 5 central mismatches to reduce binding affinity. The positive control gapmer ASO (GAP-B) targets the 64-78 region but lacks central 2'-O-methylation and hence targets 7SK for RNase H degradation. ASOs were synthesized (IDT) with phosphorothioate backbones with the following sequences:

ASO-B: mC*mC*mG*mC*mC*mU*mA*mG*mC*mC*mA*mG*mC*mC*mA*mG*mA*mU*mC
MM-B: mC*mC*mG*mC*mC*mU*mA*mC*mG*mG*mU*mC*mC*mC*mA*mG*mA*mU*mC
GAP-B: mC*mU*A*G*C*C*A*G*C*C*A*G*A*mU*mC
(m: 2'-O-Methyl RNA base; *: phosphorothioate backbone)

ASO engagement with 7SK was confirmed by DANCE-MaP experiments. 4 µg total RNA from Jurkat cells in 100 µL $H_2O$ was denatured at 98 °C for 1 min, snap cooled on 4 °C for 1 min, and then folded via addition of 100 µL of 2× folding buffer [1×: 200 mM Bicine (pH 8.0), 200 mM potassium acetate (pH 8.0) and 5 mM $MgCl_2$] (Mustoe et al., 2019) and incubated at 37 °C for 15 min. 99 µL folded RNA was then added to 1 µL of 100 µM ASO and incubated for an additional 15 min at 37 °C. Samples were then split in to two 45 µL samples and treated with DMS or ethanol as described for cell-free experiments.

HEK293T cells were seeded at 30,000 cells/well in a 96-well flat bottom plate 24 hrs prior to transfection. 100 nM of an ASO or gapmer were transfected using TransIT-Oligo (Mirus Bio). For JQ1 combination experiments, 100 nM JQ1 was added 4 hrs post transfection for a total incubation time of 20 hrs. 250 nM JQ1 was added for 24 hrs. After 24 hrs, cells were lysed in lysis buffer (Quick RNA 96-well RNA kit; Zymo) and RNA was either immediately isolated or lysed samples were flash frozen and stored at -80 °C for no longer than 48 hrs prior to RNA isolation.

**Gene expression analysis**

Total RNA was isolated (Quick RNA 96-well; Zymo) and cDNA was generated (Maxima First Strand cDNA Synthesis Kit for RT-qPCR; with dsDNase, ThermoFisher). Gene expression was assayed by RT-qPCR (using FastStart Universal SYBR Green Master; Roche) on an QuantStudio 5 instrument (Applied Biosystems). Primer sets are listed in Table S1. Primer efficiency for all targets was quantified for each run using a standard curve derived from a DNA gene fragment (gBlock; Integrated DNA Technologies; Table S1) designed to mimic the target amplicon. Expression was standardized to indicated control genes using the Pfaffl method (Michael W. Pfaffl, 2001). Data in Fig. 6D corresponds to three biological replicates from 2 independent experiments (n=6), except the 200 nM GAP-B sample which corresponds to three biological replicates from one independent experiment (n=3). Data in Fig. 6E correspond to three biological replicates from 3 independent experiments (n=9).

**Fitting adenine riboswitch titration data**

Following prior studies (Reining et al., 2013), adenine riboswitch titration data were fit assuming the three-state equilibrium

$$\mathbf{OFF} \underset{K_{pre}}{\rightleftharpoons} \mathbf{ON}_{\mathbf{-L}} \underset{K_d}{\overset{\text{ade}}{\rightleftharpoons}} \mathbf{ON}_{\mathbf{+L}}$$

where $ON_{-L}$ and $ON_{+L}$ correspond to adenine-free and adenine-bound ON states, respectively. The equilibrium constant $K_{pre}$ describes the pre-equilibrium conversion between the OFF and the ligand-free ON state, and $K_d$ is the dissociation constant of ligand binding to the ON state:

$$K_{pre} = \frac{[ON_{-L}]}{[OFF]} \quad ; \quad K_d = \frac{[ON_{-L}][ade]}{[ON_{+L}]}$$

The total RNA concentration [RNA] is the sum of the three species:

$$[RNA] = [OFF] + [ON_{-L}] + [ON_{+L}] = [OFF] + K_{pre}\,[OFF] + \frac{K_{pre}}{K_d}[OFF][ade] \tag{3}$$

The amount of free ligand, [ade], can be further expressed as a function of the total ligand $[ade_0]$:

$$[ade] = [ade_0] - [ON_{+L}] = [ade_0] - [RNA] + [OFF] + K_{pre}\,[OFF] \tag{4}$$

Substituting (3) into (4) and solving for [OFF] (Mathematica) yields two solutions, the valid one is:

$$[OFF] = \frac{\sqrt{4\,K_d\left(K_{pre} + K_{pre}^2\right)[RNA] + \left(K_d + K_d\,K_{pre} + K_{pre}[ade_0] - K_{pre}[RNA]\right)^2}}{2\,K_{pre}\left(K_{pre} + 1\right)} + \frac{-K_d - K_d\,K_{pre} - K_{pre}[ade_0] + K_{pre}[RNA]}{2\,K_{pre}\left(K_{pre} + 1\right)}$$

The combined fraction of species in the ON state, $f_{ON}$, is then:

$$f_{ON} = 1 - f_{alt} = 1 - \frac{[OFF]}{[RNA]}$$

To account for incomplete saturation, we fit to the equation:

$$f_{ON} = m * \left(1 - \frac{[OFF]}{[RNA]}\right)$$

where $m$ is the fractional saturation, [*RNA*] and [$ade_0$] are the input concentrations, and $f_{ON}$ is the population measured by DANCE-MaP. Fits were obtained using the curve_fit module of SciPy in Python. The modest deviations of the data from the expected curve are likely to due to DMS modification of the adenine ligand, which reduces effective ligand concentration, and a central tendency bias of DANCE-MaP for clustering into more-equally-weighted groups (for example, a 75:25 ratio may be resolved as a 70:30 ratio). The fitted saturation fraction (*m*) values were 0.77 for both replicates, consistent with prior studies (Warhaut et al., 2017).

**Data and code availability**

Structure probing data have been deposited in the NCBI BioProject database (http://www.ncbi.nlm.nih.gov/bioproject/741330) and will be released upon publication. The fully automated *DanceMapper* pipeline was written in Python (2.7) and Cython. *DanceMapper*, along with scripts for performing structure modeling and visualizing DANCE solutions, will be available for download at https://github.com/MustoeLab/DanceMapper.

**Supporting Figure Legends**

**Figure S1:** Schematic of the DANCE-MaP algorithm. In read diagrams at top, 0 indicates no mutation (match), 1 indicates a mutation, and (*) indicates no data. See Methods for further details.

**Figure S2:** Benchmarking DANCE ML deconvolution on synthetic data. Ensembles consisting of 2, 3, 4, and 5 distinct reactivity profiles were generated for a 200 nt "RNA". Profiles were constructed by randomly drawing reactivities from *E. coli* 16S and 23S rRNA cell-free DMS-MaP modification datasets (Mustoe et al., 2019) and systematically varying the fraction of nucleotides that differ between each profile in the ensemble (*top*). The population splits of each ensemble were also systematically varied. These reference ensembles were then used to generate 500,000 synthetic reads, with 10% of positions in each read randomly assigned as missing data, and 20% of nucleotides randomly "inactivated" (representing worse-than-typical data quality). DANCE-MaP was then used to deconvolute the synthetic data. Each colored square represents the difference between the ML solution and the known generating reference, as a function of population ($\pi$) and reactivity ($\mu$) differences (see key). When DANCE-MaP yielded an incomplete deconvolution (for example, predicted a 2-state ensemble when the reference ensemble had 3 states), the difference is computed using the most similar substates of the reference and is marked with diagonal hatching. Low quality solutions (defined in Methods) are indicated by dense plaid hatching. Ensembles for which DANCE-MaP was unable to detect any heterogeneity (yielding a 1-component ensemble) are white. Data are presented from left-to-right for increasing difference between ensemble profiles, and bottom-to-top for increasing minimum population. Nucleotide positions were considered to be different if their reactivity rates differed by >0.01. Accurate deconvolution depends on both extent of structural differences between ensemble states and their relative populations. For low population states, or for ensembles of consisting of very similar states, deconvolution is incomplete but $\mu$ errors are low, with the minor state being absorbed into more populous states. Deconvolution performance decreases for heterogenous ensembles consisting of 4-5 highly different states. This analysis likely overestimates errors because reference ensembles were randomly constructed, and highly divergent states are more likely to contain locally anticorrelated reactivities which are rejected as artifactual by DANCE-MaP.

**Figure S3:** Structural ensembles of native sequence *add* riboswitch constructs, as a function of adenine concentration. In each structure diagram, pairing probabilities are shown at top, deconvoluted DMS reactivities at middle, and deconvoluted PAIRs at bottom. The ligand-free and adenine-bound ON states are resolved as a single ON state due to the small number of nucleotides that change reactivity upon ligand binding. In one exception, three states corresponding to ON (ligand-bound), OFF, and ON (ligand-free) were resolved at 6.2 μM adenine concentration (third state not shown). Pairing probabilities and PAIRs were reproducible across full independent replicate titrations (not shown).

**Figure S4:** Structural ensembles for the $Mut_{ON}$ and $Mut_{OFF}$ *add* riboswitch mutants, measured in the presence of 0 and 500 μM adenine. In each structure diagram, pairing probabilities are shown at top, deconvoluted DMS reactivities at middle, and deconvoluted PAIRs at bottom.

**Figure S5:** Benchmarking DANCE PAIR and RING deconvolution on synthetic data. Shown are results from measuring correlations for 1-nt windows (RING analysis); comparable results were obtained when measuring correlations over 3-nt windows (PAIR analysis). 300 3-state ensembles of varying populations and reactivity heterogeneity were constructed for a 200 nt “RNA”. For each ensemble state, pairs of nucleotides were assigned to undergo correlated modification. State reactivities and correlations were drawn from *E. coli* 16S and 23S rRNA cell-free DMS modification datasets (Mustoe et al., 2019). Each reference ensemble was then used to generate 1,500,000 synthetic reads, with 10% of each read randomly set as missing data. Synthetic reads were deconvoluted and assigned using *DanceMapper*, and RING analysis was performed on each state. Positive predictive value (ppv) and sensitivity (sens) of the measured RINGs were then computed in relation to correlations associated with each reference state. When DANCE yielded an incomplete deconvolution (for example, predicted a 2-state ensemble), ppv/sens are computed to the most similar reference state and are marked with horizontal hatching. Automatically detected low quality deconvolutions (as defined in Methods) are indicated by plaid hatching. White is used to indicate that a state was not detected (incomplete deconvolution) or no RINGs were detected. For fully deconvoluted solutions, ppv>0.95 (mean=0.999). ppv is lower for incompletely deconvoluted solutions, reflecting the presence of indirect correlations arising from state heterogeneity and direct correlations of “absorbed” states. sens also decreases for incompletely deconvoluted solutions because noise from state heterogeneity conceals weak correlations. In addition to state purity, sens depends strongly on the number of reads assigned to each state. Hence, sens is best for the most populous state (which has the most reads in the parent dataset), and decreases predictably for less populous states.

**Figure S6:** State-specific PAIR and RING correlations measured for 7SK RNA. (A) PAIR and RING correlations for in-cell and cell-free RNA measured from a single deeply sequenced sample (same as shown in Figure 3) and a consolidated replicate constructed by pooling multiple lower depth independent replicates. Minimum free energy structures are shown as arcs (top, gray). High- and moderate-confidence PAIRs are shown as dark and light blue arcs, respectively. Through-space RING correlations are shown as high (dark red, G>100) and moderate (light red, G>20) confidence arcs. RINGs were filtered for contact distance (>15) (Hajdin et al., 2013; Dethoff et al., 2018), and only positive correlations are shown (Mustoe et al., 2019). (B) RING correlations from the deeply sequenced sample superimposed on the consensus models for states A and B. In addition to state-specific contact distance filtering used in A, RINGs were additionally filtered by contact distance (<15) in relation to the other structural state to exclude any residual signals originating from the alternative secondary structure. State B data are reproduced from Figure 3D.

**Figure S7:** Comparison of states A and B, defined by DANCE-MaP, with prior models of the 7SK RNA structure (Wassarman and Steitz, 1991; Marz et al., 2009; Brogie and Price, 2017). (A) DANCE-MaP consensus secondary structures models for states A and B. Base pairs are shown as arcs; structural landmarks are labeled for each state. (B-D) Comparison of DANCE-deconvoluted states A and B with previously described models for the 7SK RNA. Base pairs shared between DANCE-deconvoluted and prior models, unique to DANCE models, and unique to prior models are gray, green and blue, respectively. The Brogie and Price high $Mg^{2+}$ model (panel D) was inferred using the partial structure published in Figure 2B, and using the reactivities supplied in the supplementary information to model remaining positions (Brogie and Price, 2017).

**Figure S8:** Determination of consensus 7SK RNA structures. (A) Expanded view of DMS reactivities for the SL0 region for states A and B, in cells. The state A inset shows the alternative SL1ext pairing. Green and purple boxes indicate nucleotides paired in SL0 and SL1ext, respectively. For state B, reactivities and PAIRs supporting formation of an extension of SL0 and SL2ext are shown. (B, C) Comparisons of in-cell and cell-free structures modeled with and without PAIR data. The SL2ext region, which is poorly defined in the absence of PAIR data, is highlighted.

**Figure S9:** DANCE-MaP characterization of *in vitro* transcribed native sequence and mutant 7SK RNAs. (A) Comparison of DMS reactivities for states A and B for native *in vitro* transcribed versus cell-free 7SK RNA. Pearson's R is shown. (B) DANCE-MaP-derived structural ensembles for each *in vitro* RNA. States were assigned to A/A-like and B/B-like based on comparison to the native sequence RNA. States were assigned by the following criteria: has clear SL1: A or A-like; has clear SL1alt and/or SL2ext: B or B-like. Percentages assigned to each state are shown. Data are representative of three (native, M1, M1+M2, M3) or two (M4) independent replicates. Populations denote means and standard deviations across replicates. Note that observed PAIRs are limited due to low sequencing depth.

**Figure S10:** Structural models for deconvoluted 7SK states in perturbed (A) Jurkat and (B) RPE-1 cells. Proliferating Jurkat or RPE-1 cells were treated with vehicle (0.01% DMSO) or 1 μM flavopiridol. RPE-1 quiescence was established by contact inhibition. Data are representative of at least two independent replicates. In-cell Jurkat and proliferating RPE-1 data are provided for reference and are reproduced from Figures 3 and 5.

**Table S1: Primer and template sequences**

| | |
|---|---|
| Primers | |
| 7SK-RT | AAAAGAAAGGCAGACTGCCAC |
| 7SK-PCR1-F | GACTGGAGTTCAGACGTGTGCTCTTCCGATCTNNNNNGGATGTGAGGGCGATCTG |
| 7SK-PCR1-R | CCCTACACGACGCTCTTCCGATCTNNNNNAAAAGAAAGGCAGACTGCCACATG |
| PCR gBlock template primers | |
| 7SK-Template-F | GAAATTAATACGACTCACTATAGGGATGTGAG |
| 7SK-Template-R | AmAAAGAAAGGCAGACTGCCAC<br>mA-2'-OMe Adenosine |
| g-blocks for IVT | |
| *add* riboswitch Mut$_{ON}$ | TAATACGACTCACTATAGGCCTTCGGGCCAAGATCAACGCTTCATATAATCCTCGTGATATGGTCGGGGAGTTTCTACCAAGAGCCTTAAACTCTTGATTATGAAGTCTGTCGCTTTATCCGAAATTTTATAAAGAGAAGACTCATGAATTCGATCCGGTTCGCCGGATCCAAATCGGGCTTCGGTCCGGTTC |
| *add* riboswitch Mut$_{OFF}$ | TAATACGACTCACTATAGGCCTTCGGGCCAAGATCAACGCTTCATATAATCCTAATGATATGGTTTGGAAGTTTCTACCAAGAGCCTTAAACTCTTGATTATGAAGTCTGTCGCTTTATCCGAAATTTTATAAAGAGAAGACTCATGAATTCGATCCGGTTCGCCGGATCCAAATCGGGCTTCGGTCCGGTTC |
| Native 7SK sequence | GAAATTAATACGACTCACTATAGGGATGTGAGGGCGATCTGGCTGCGACATCTGTCACCCCATTGATCGCCAGGGTTGATTCGGCTGATCTGGCTGGCTAGGCGGGTGTCCCCTTCCTCCCTCACCGCTCCATGTGCGTCCCTCCCGAAGCTGCGCGCTCGGTCGAAGAGGACGACCATCCCCGATAGAGGAGGACCGGTCTTCGGTCAAGGGTATACGAGTAGCTGCGCTCCCCTGCTAGAACCTCCAAACAAGCTCTCAAGGTCCATTTGTAGGAGAACGTAGGGTAGTCAAGCTTCCAAGACTCCAGACACATCCAAATGAGGCGCTGCATGTGGCAGTCTGCCTTTCTTTT |

| | |
|---|---|
| 7SK M1 | GAAATTAATACGACTCACTATAGGGATGTGAGGGCGATCTG<br>GCTGCGAC**TAG**TGTCACCCCATTGATCGCCAGGGTTGATT<br>CGGCTGATCTGGCTGGCTAGGCGGGTGTCCCCTTCCTCCC<br>TCACCGCTCCATGTGCGTCCCTCCCGAAGCTGCGCGCTCG<br>GTCGAAGAGGACGACCATCCCCGATAGAGGAGGACCGGT<br>CTTCGGTCAAGGGTATACGAGTAGCTGCGCTCCCCTGCTA<br>GAACCTCCAAACAAGCTCTCAAGGTCCATTTGTAGGAGAAC<br>GTAGGGTAGTCAAGCTTCCAAGACTCCAGACACATCCAAAT<br>GAGGCGCTGCATGTGGCAGTCTGCCTTTCTTTT |
| 7SK M1+M2 | GAAATTAATACGACTCACTATAGGGATGTGAGGGCGATCTG<br>GCTGCGAC**TAG**TGTCACCCCATTGATCGCCAGGGTTGATT<br>CGGCTGATCTGGCTGGCTAGGCG**CTA**GTCCCCTTCCTCCC<br>TCACCGCTCCATGTGCGTCCCTCCCGAAGCTGCGCGCTCG<br>GTCGAAGAGGACGACCATCCCCGATAGAGGAGGACCGGT<br>CTTCGGTCAAGGGTATACGAGTAGCTGCGCTCCCCTGCTA<br>GAACCTCCAAACAAGCTCTCAAGGTCCATTTGTAGGAGAAC<br>GTAGGGTAGTCAAGCTTCCAAGACTCCAGACACATCCAAAT<br>GAGGCGCTGCATGTGGCAGTCTGCCTTTCTTTT |
| 7SK M3 | GAAATTAATACGACTCACTATAGGGATGTGAGGGCGATCTG<br>GCTGCGACATCTGTCACCCCATTGATCGCCAGGGTTGATT<br>CGGCTGATCTGGCTGGCTAGGCGGGTGTCCCCTTCCTCCC<br>TCACCGCTCCATGTGCGTCCCTCCCGAAGCTGCGCGCTCG<br>GTCGAAGAGGACGACCATCCCCGATAGAGGAGGACCGGT<br>CTTCGGTCAAGGGTATACGAGTAGCTGCGCTCCCCTGCTA<br>GAACCTCCAAACAAGCTCTCAAGGTCCATTTGTAGGAGAAC<br>GTAGGGTAGTCAAGCTTCCAAGACTCCAGA**ATGTAAA**AAAT<br>GAGGCGCTGCATGTGGCAGTCTGCCTTTCTTTT |
| 7SK M4 | GAAATTAATACGACTCACTATAGGGATGTGAGGGCGATCTG<br>GCTGCGACATCTGTCACCCCATTGATCGCCAGGGTTGATT<br>CGGCTGATCTGGCTGGCTAGGCGGGTGTCCC**TAC**CCTCCC<br>TCACCGCTCCATGTGCGTCCCTCCCGAAGCTGCGCGCTCG<br>GTCGAAGAGGACGACCATCCCCGATAGAGGAGGACCGGT<br>CTTCGGTCAAGGGTATACGAGTAGCTGCGCTCCCCTGCTA<br>GAACCTCCAAACAAGCTCTCAAGGTCCATTTGTAGGAGAAC |

| | |
|---|---|
| | GTAGGGTAGTCAAGCTTCCAAGACTCCAGACACATCCAAAT GAGGCGCTG<u>CATGTGGCAGTCTGCCTTTCTTTT</u> |
| Mutations are in **bold**<br>Primer binding regions are <u>underlined</u> | |
| qPCR primers | |
| 7SK-F | CCTGCTAGAACCTCCAAACAA |
| 7SK-R | GGAGTCTTGGAAGCTTGACTAC |
| HEXIM1-F | CCGAGGCCAGTAAGTTGGG |
| HEXIM1-R | GACGGGCGTCTCCTATGTTT |
| TBP-F | GAGAGTTCTGGGATTGTACCG |
| TBP-R | ATCCTCATGATTACCGCAGC |
| RPL13a-F | GCCTACAAGAAAGTTTGCCTATC |
| RPL13a-R | TGGCTTTCTCTTTCCTCTTCTC |
| GAPDH-F | GTCAACGGATTTGGTCGTATTG |
| GAPDH-R | TGTAGTTGAGGTCAATGAAGGG |
| gBlocks for qPCR standardization | |
| 7SK-qPCR | GGATGTGAGGGCGATCTGGCTGCGACATCTGTCACCCCAT TGATCGCCAGGGTTGATTCGGCTGATCTGGCTGGCTAGGC GGGTGTCCCCTTCCTCCCTCACCGCTCCATGTGCGTCCCT CCCGAAGCTGCGCGCTCGGTCGAAGAGGACGACCATCCC CGATAGAGGAGGACCGGTCTTCGGTCAAGGGTATACGAGT AGCTGCGCTCCCCTGCTAGAACCTCCAAACAAGCTCTCAA GGTCCATTTGTAGGAGAACGTAGGGTAGTCAAGCTTCCAA GACTCCAGACACATCCAAATGAGGCGCTGCATGTGGCAGT CTGCCTTTCTTTT |
| HEXIM-qPCR | AGCCTTGTCATGACTCCGAGGCCAGTAAGTTGGGGGCTCC TGCCGCAGGGGGCGAAGAGGAGTGGGGACAGCAGCAGAG ACAGCTGGGGAAGAAAAAACATAGGAGACGCCCGTCCAAG AAGAAGC |
| TBP-qPCR | GCCAGCTTCGGAGAGTTCTGGGATTGTACCGCAGCTGCAA AATATTGTATCCACAGTGAATCTTGGTTGTAAACTTGACCTA AAGACCATTGCACTTCGTGCCCGAAACGCCGAATATAATCC CAAGCGGTTTGCTGCGGTAATCATGAGGATAAGAGAGCCA |

| | |
|---|---|
| RPL13a-qPCR | ATCCCACCGCCCTACGACAAGAAAAAGCGGATGGTGGTTC CTGCTGCCCTCAAGGTCGTGCGTCTGAAGCCTACAAGAAA GTTTGCCTATCTGGGGCGCCTGGCTCACGAGGTTGGCTGG AAGTACCAGGCAGTGACAGCCACCCTGGAGGAGAAGAGG AAAGAGAAAGCCAAGATCCACTACCGGAAGAAGAAACAGC TCATGAGGCTACGGAAACAGGCCGAGAAGAACGTGGAGAA GAAAATTGACAAATACACAGAGGTCCTCAAGACCCACGGA CTCCTGGTC |
| GAPDH-qPCR | GAAGGTCGGAGTCAACGGATTTGGTCGTATTGGGCGCCTG GTCACCAGGGCTGCTTTTAACTCTGGTAAAGTGGATATTGT TGCCATCAATGACCCCTTCATTGACCTCAACTACATGGTTT ACATGTTCCAATATGATTCCACCCATGGCAAATTCCATGGC ACCGTCAAGGCTGAGAACGGGAAGCTTGTCATCAATGGAA AT |

### Figure S1

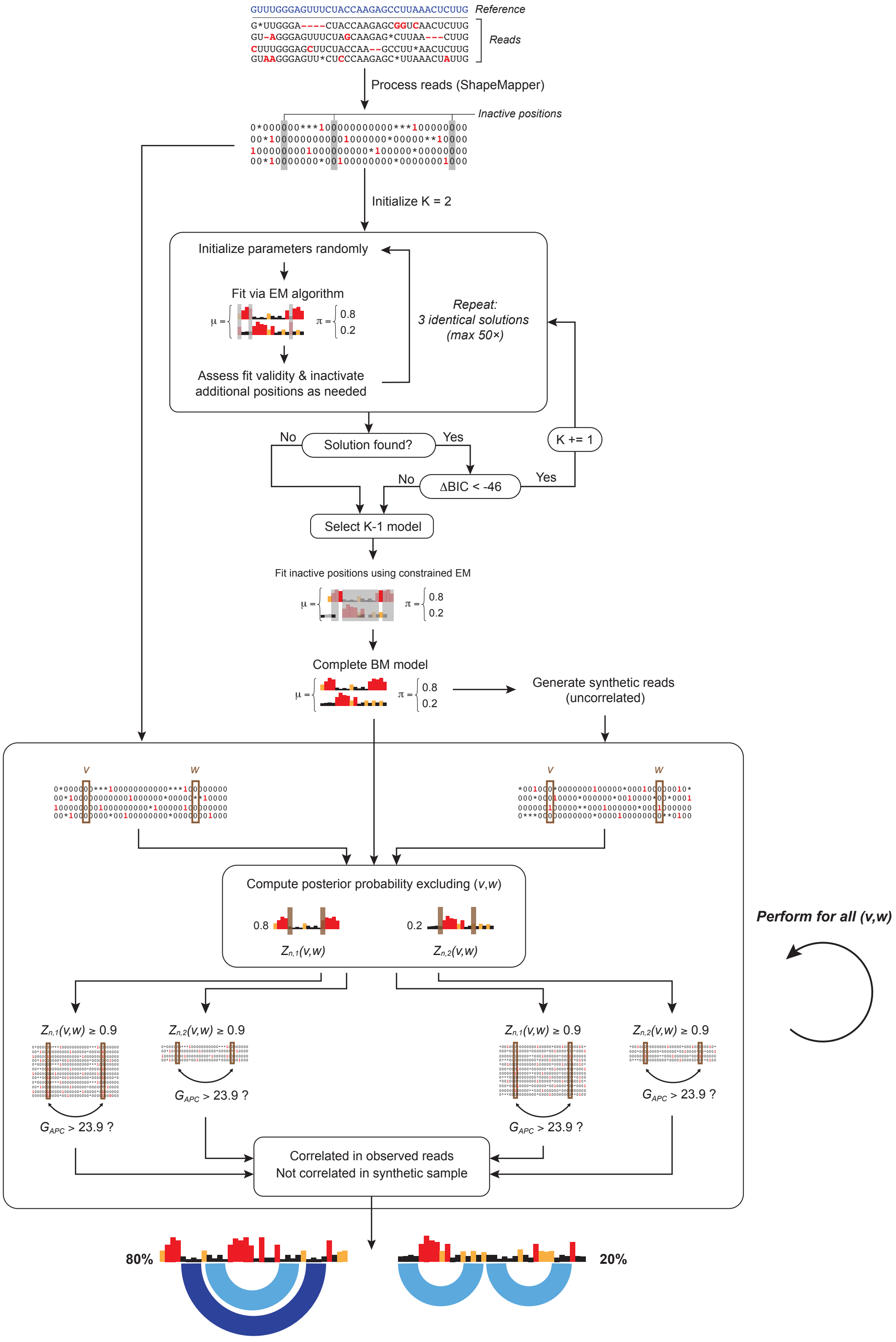

GUUUGGGAGUUUCUACCAAGAGCCUUAAACUCUUG Reference
Reads
Process reads (ShapeMapper)
Inactive positions
Initialize K = 2
Initialize parameters randomly
Fit via EM algorithm
Assess fit validity & inactivate additional positions as needed
Repeat: 3 identical solutions (max 50×)
Solution found?
No
Yes
ΔBIC < -46
K += 1
Select K-1 model
Fit inactive positions using constrained EM
Complete BM model
Generate synthetic reads (uncorrelated)
Compute posterior probability excluding (v,w)
Perform for all (v,w)
Correlated in observed reads
Not correlated in synthetic sample
80%
20%

### Figure S2

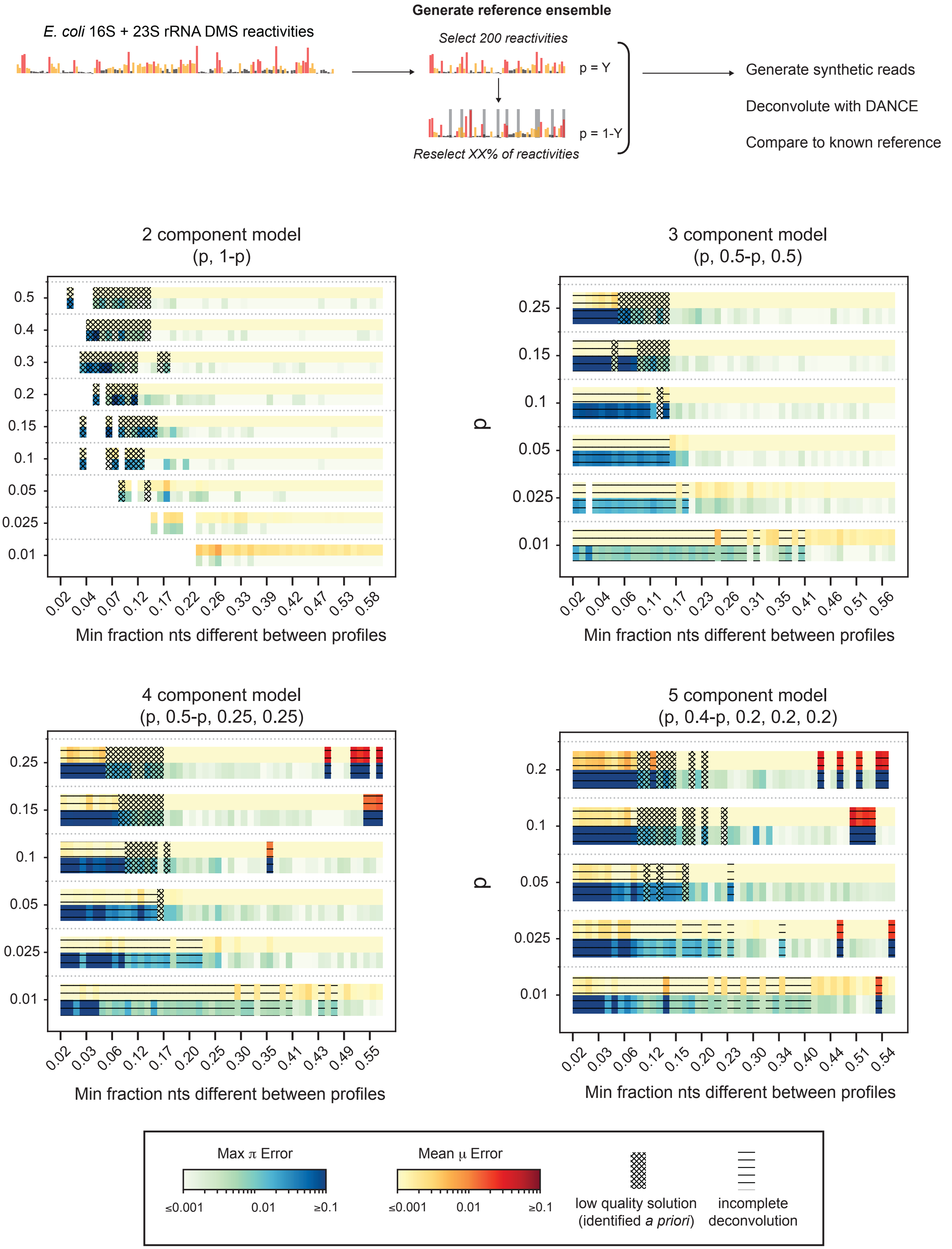

E. coli 16S + 23S rRNA DMS reactivities
Generate reference ensemble
Select 200 reactivities
p = Y
p = 1-Y
Reselect XX% of reactivities
Generate synthetic reads
Deconvolute with DANCE
Compare to known reference
2 component model
(p, 1-p)
p
0.5
0.4
0.3
0.2
0.15
0.1
0.05
0.025
0.01
0.02 0.04 0.07 0.12 0.17 0.22 0.26 0.33 0.39 0.42 0.47 0.53 0.58
Min fraction nts different between profiles
3 component model
(p, 0.5-p, 0.5)
p
0.25
0.15
0.1
0.05
0.025
0.01
0.02 0.04 0.06 0.11 0.17 0.23 0.26 0.31 0.35 0.41 0.46 0.51 0.56
Min fraction nts different between profiles
4 component model
(p, 0.5-p, 0.25, 0.25)
p
0.25
0.15
0.1
0.05
0.025
0.01
0.02 0.03 0.06 0.12 0.17 0.20 0.25 0.30 0.35 0.41 0.43 0.49 0.55
Min fraction nts different between profiles
5 component model
(p, 0.4-p, 0.2, 0.2, 0.2)
p
0.2
0.1
0.05
0.025
0.01
0.02 0.03 0.06 0.12 0.15 0.20 0.23 0.30 0.34 0.40 0.44 0.51 0.54
Min fraction nts different between profiles
Max π Error
≤0.001 0.01 ≥0.1
Mean μ Error
≤0.001 0.01 ≥0.1
low quality solution (identified a priori)
incomplete deconvolution

### Figure S3

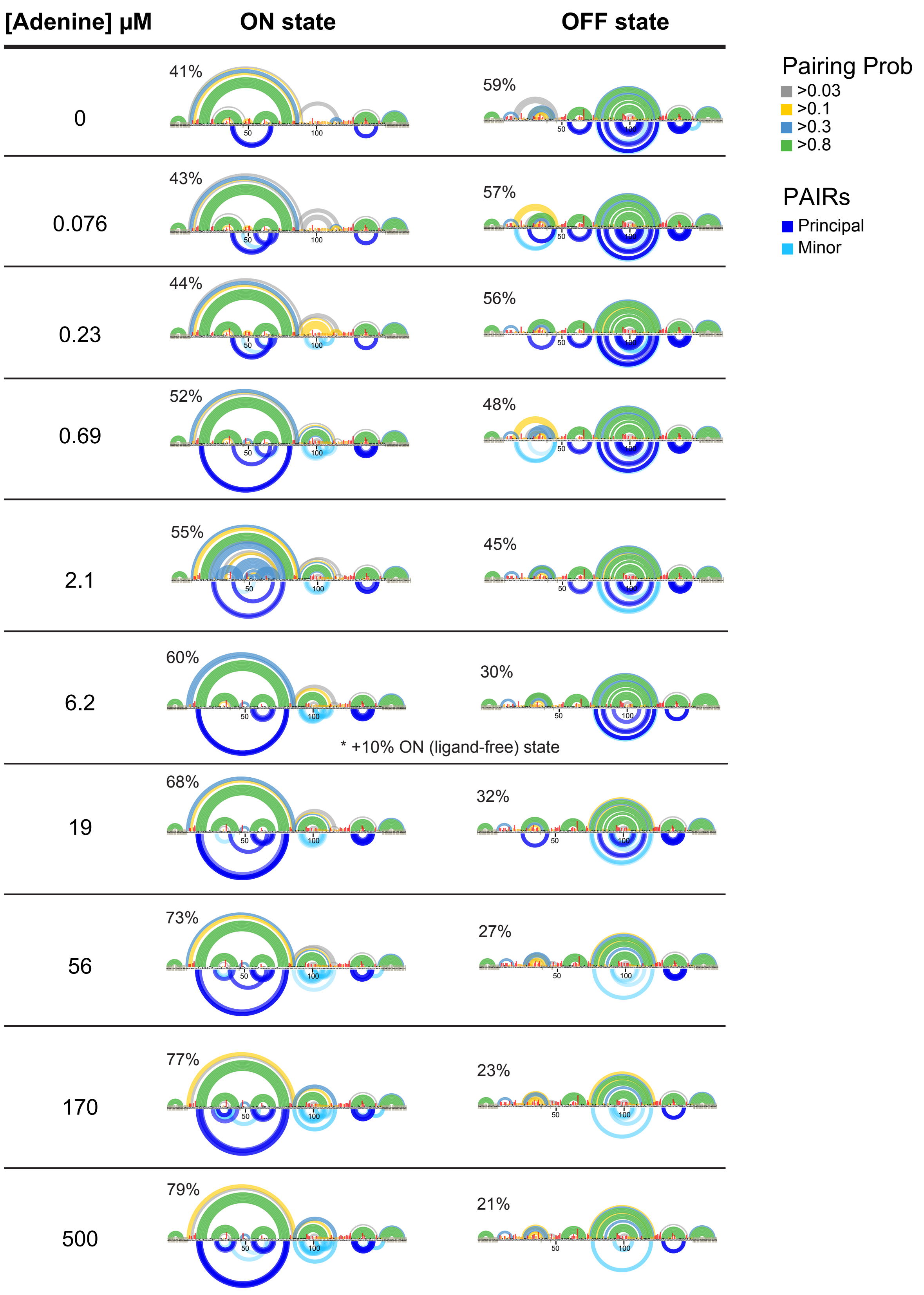

[Adenine] µM
ON state
OFF state
0
41%
59%
0.076
43%
57%
0.23
44%
56%
0.69
52%
48%
2.1
55%
45%
6.2
60%
30%
* +10% ON (ligand-free) state
19
68%
32%
56
73%
27%
170
77%
23%
500
79%
21%
Pairing Prob
>0.03
>0.1
>0.3
>0.8
PAIRs
Principal
Minor

### Figure S4

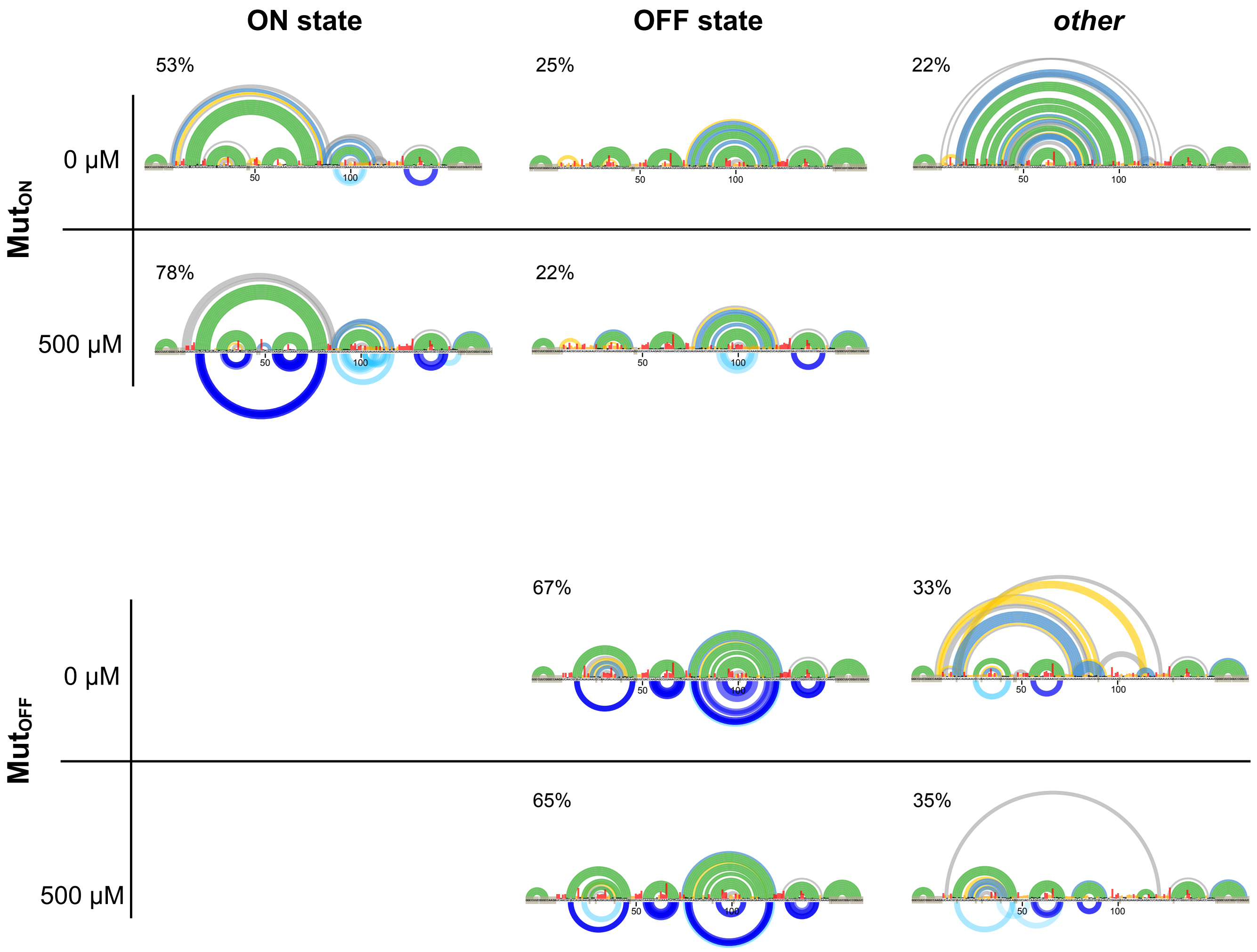

ON state
OFF state
other
MutON
0 μM
500 μM
53%
25%
22%
78%
22%
MutOFF
0 μM
500 μM
67%
33%
65%
35%
50
100

### Figure S5

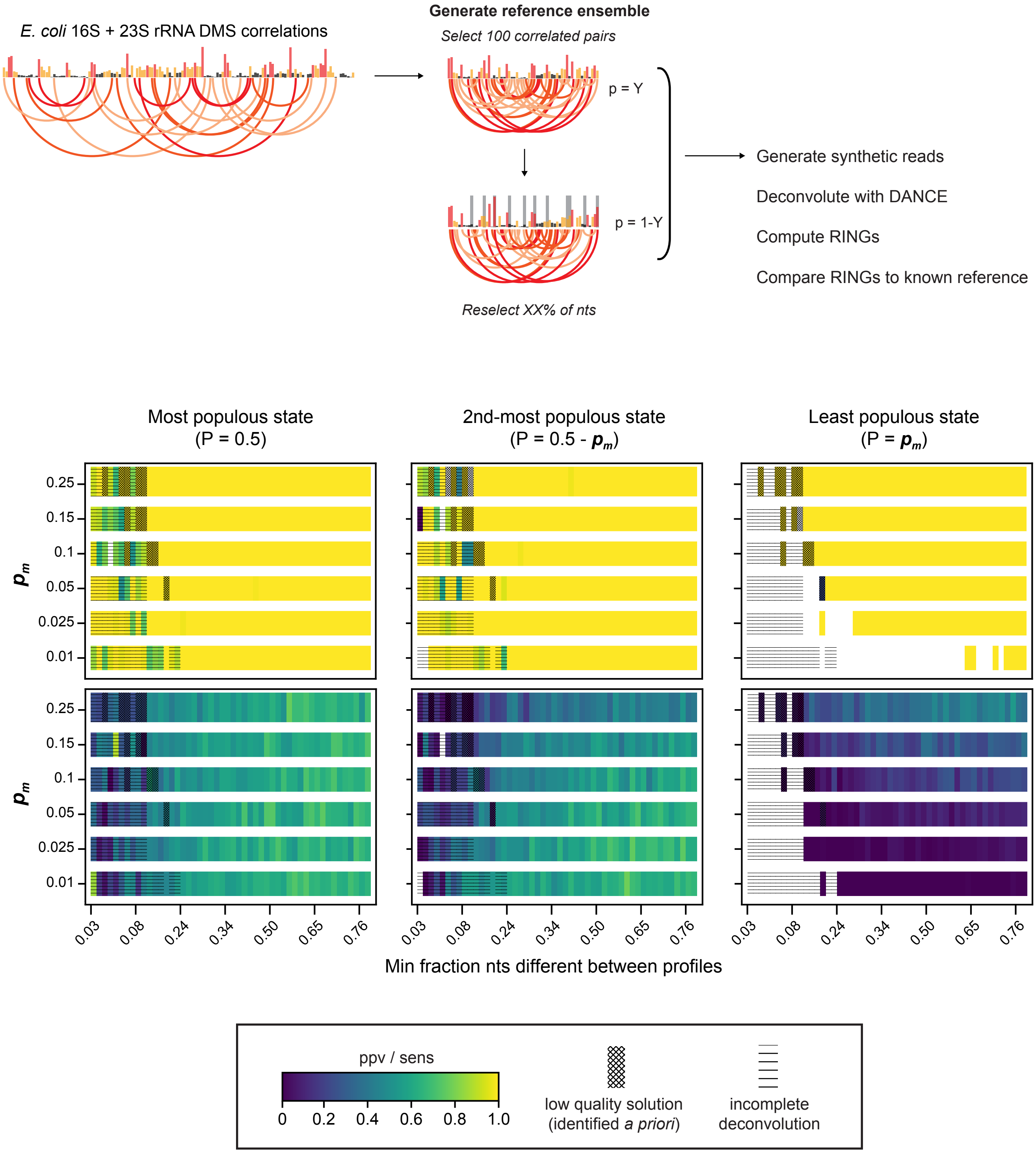

E. coli 16S + 23S rRNA DMS correlations
Generate reference ensemble
Select 100 correlated pairs
p = Y
p = 1-Y
Reselect XX% of nts
Generate synthetic reads
Deconvolute with DANCE
Compute RINGs
Compare RINGs to known reference
Most populous state
(P = 0.5)
2nd-most populous state
(P = 0.5 - $p_m$)
Least populous state
(P = $p_m$)
ppv
sens
$p_m$
0.25
0.15
0.1
0.05
0.025
0.01
0.03
0.08
0.24
0.34
0.50
0.65
0.76
Min fraction nts different between profiles
ppv / sens
0
0.2
0.4
0.6
0.8
1.0
low quality solution
(identified a priori)
incomplete
deconvolution

Figure S6

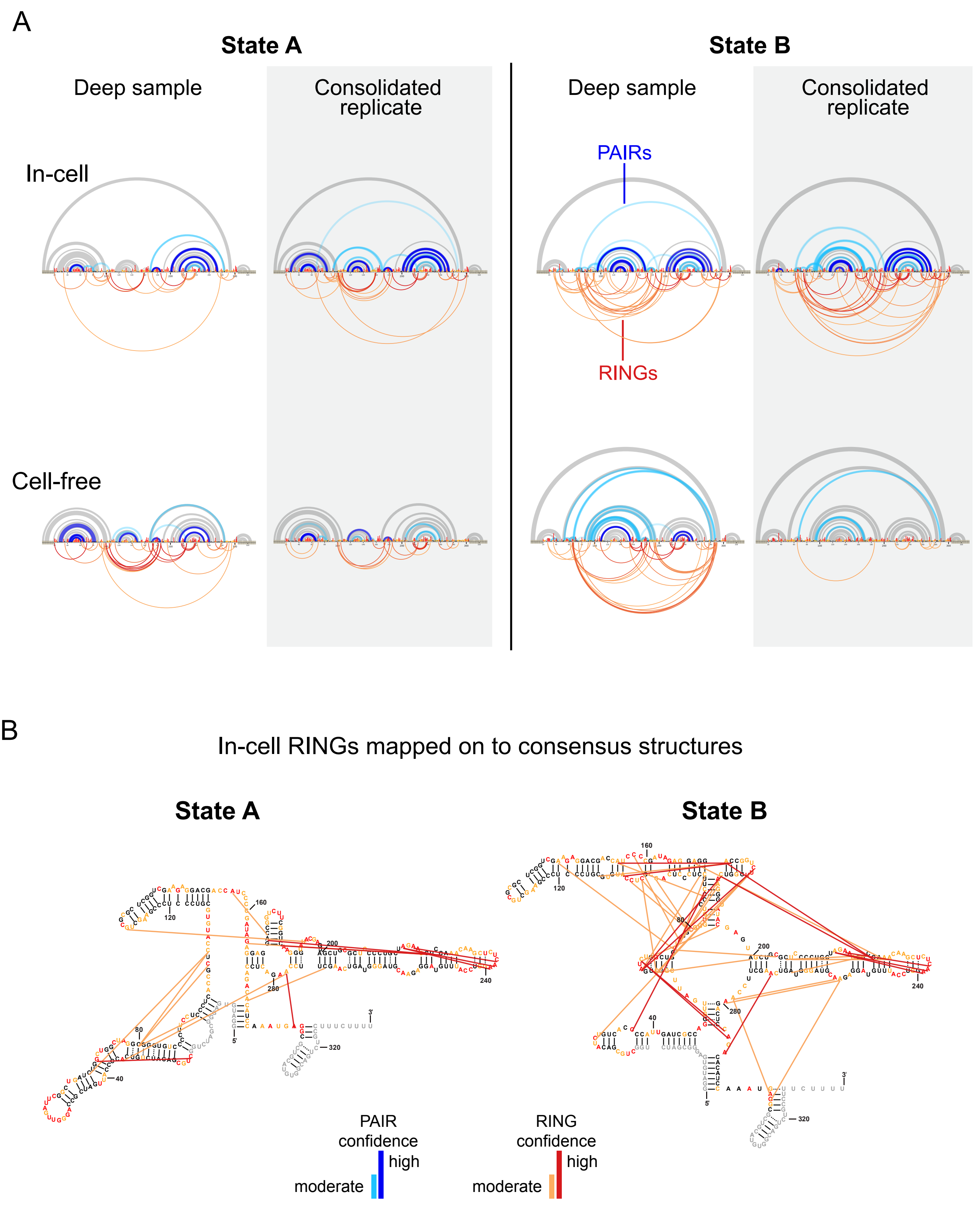

A
State A
State B
Deep sample
Consolidated replicate
Deep sample
Consolidated replicate
In-cell
PAIRs
RINGs
Cell-free
B
In-cell RINGs mapped on to consensus structures
State A
State B
PAIR confidence
high
moderate
RING confidence
high
moderate

Figure S7

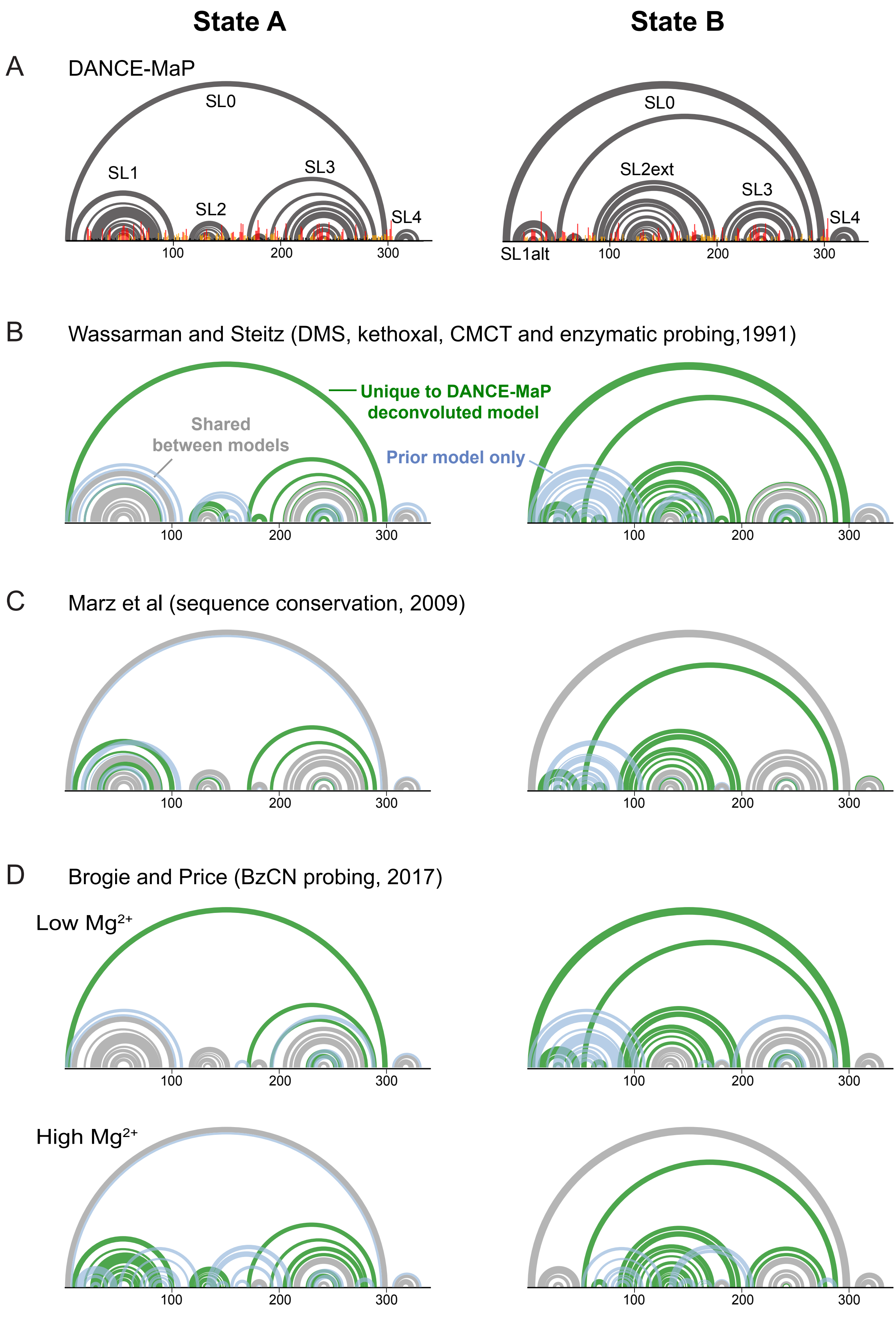

State A
State B
A
DANCE-MaP
SL0
SL1
SL2
SL3
SL4
100
200
300
SL0
SL2ext
SL3
SL4
SL1alt
100
200
300
B
Wassarman and Steitz (DMS, kethoxal, CMCT and enzymatic probing,1991)
Unique to DANCE-MaP deconvoluted model
Shared between models
Prior model only
100
200
300
100
200
300
C
Marz et al (sequence conservation, 2009)
100
200
300
100
200
300
D
Brogie and Price (BzCN probing, 2017)
Low Mg2+
100
200
300
100
200
300
High Mg2+
100
200
300
100
200
300

### Figure S8

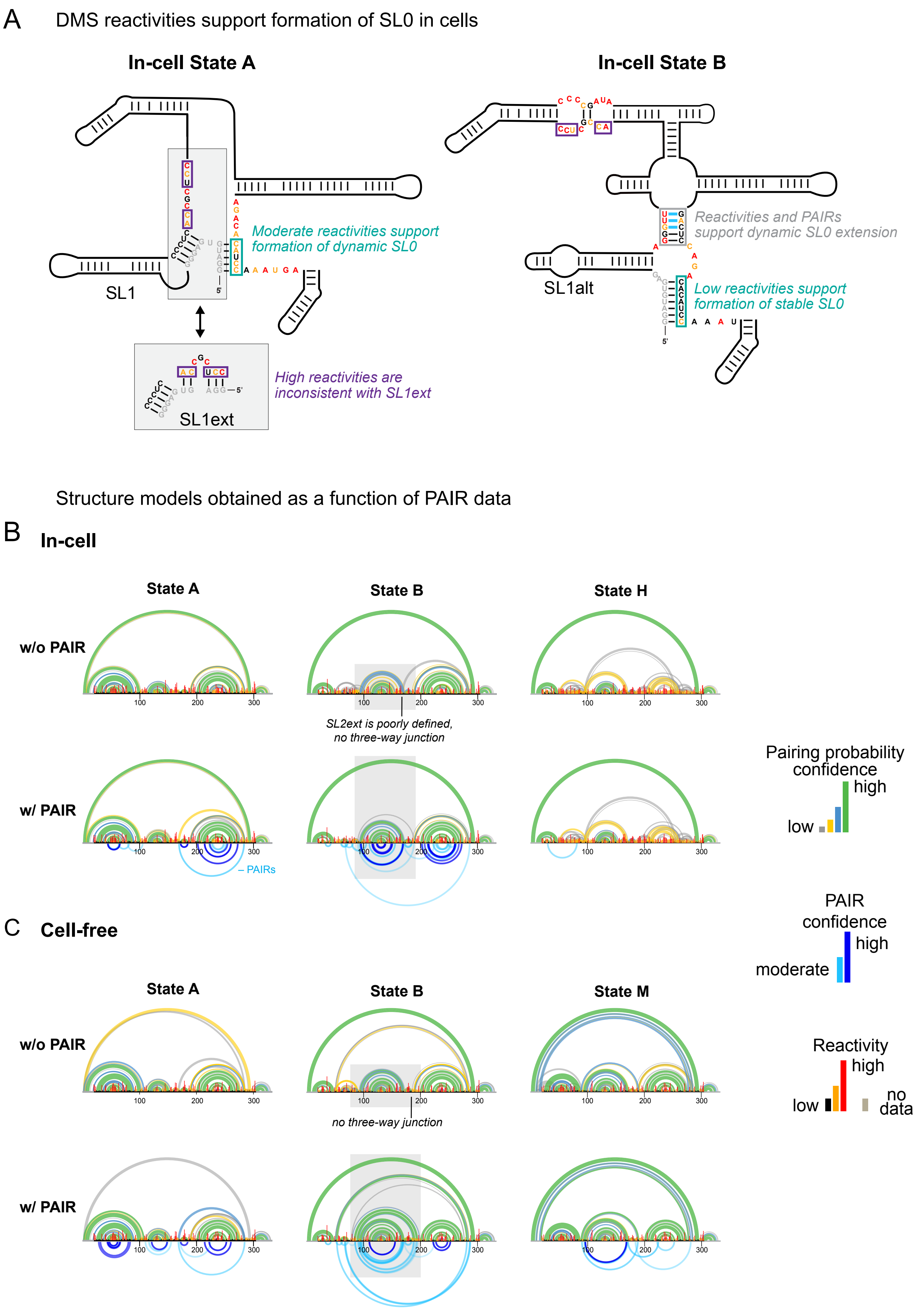
A
DMS reactivities support formation of SL0 in cells
In-cell State A
In-cell State B
Moderate reactivities support formation of dynamic SL0
SL1
SL1ext
High reactivities are inconsistent with SL1ext
Reactivities and PAIRs support dynamic SL0 extension
SL1alt
Low reactivities support formation of stable SL0
Structure models obtained as a function of PAIR data
B
In-cell
State A
State B
State H
w/o PAIR
w/ PAIR
SL2ext is poorly defined, no three-way junction
– PAIRs
Pairing probability confidence
high
low
PAIR confidence
high
moderate
C
Cell-free
State A
State B
State M
w/o PAIR
w/ PAIR
no three-way junction
Reactivity
high
low
no data

### Figure S9

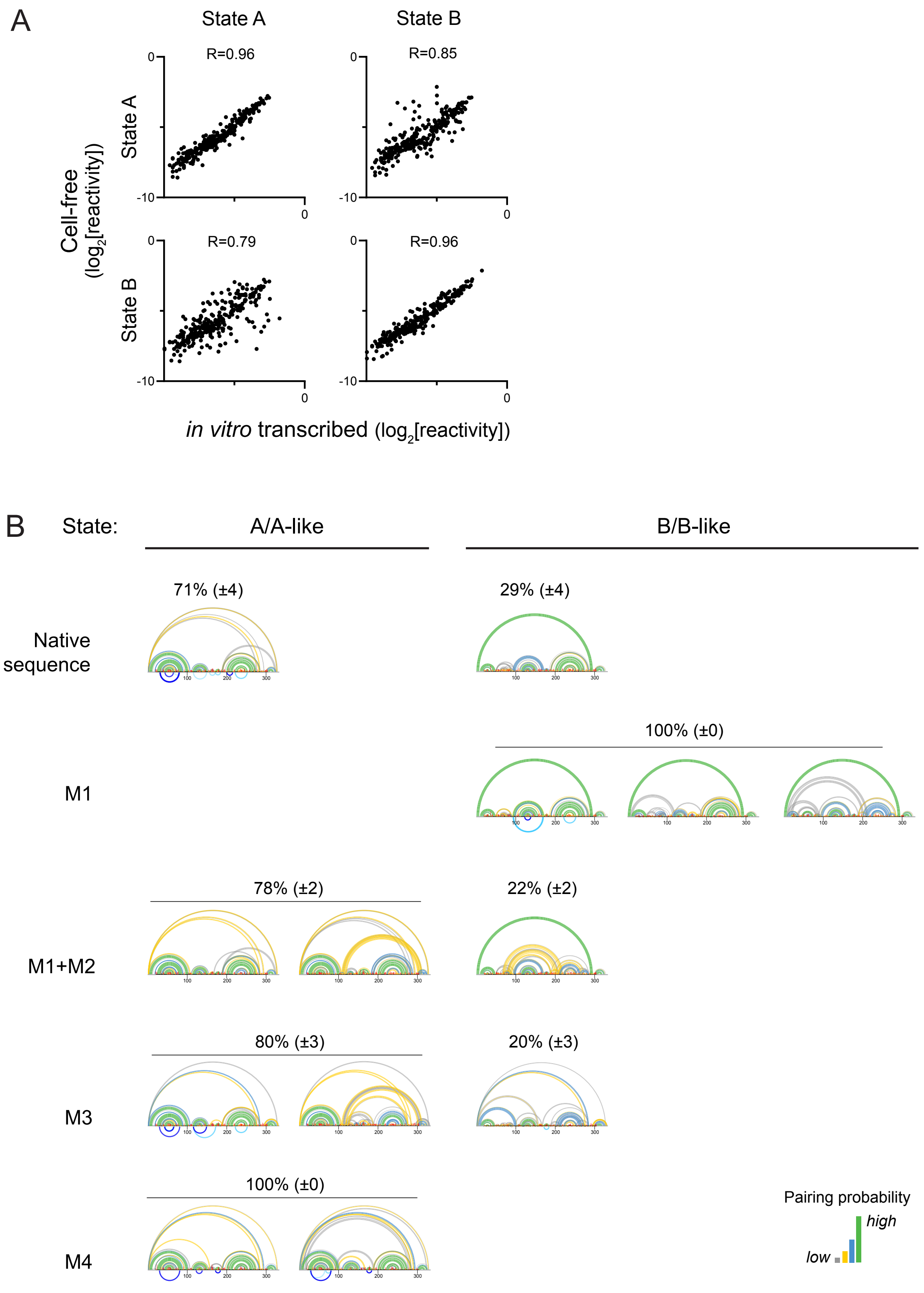
A
State A
State B
R=0.96
R=0.85
R=0.79
R=0.96
Cell-free (log2[reactivity])
State A
State B
0
-10
in vitro transcribed (log2[reactivity])
B
State:
A/A-like
B/B-like
71% (±4)
29% (±4)
Native sequence
100% (±0)
M1
78% (±2)
22% (±2)
M1+M2
80% (±3)
20% (±3)
M3
100% (±0)
M4
Pairing probability
high
low

### Figure S10

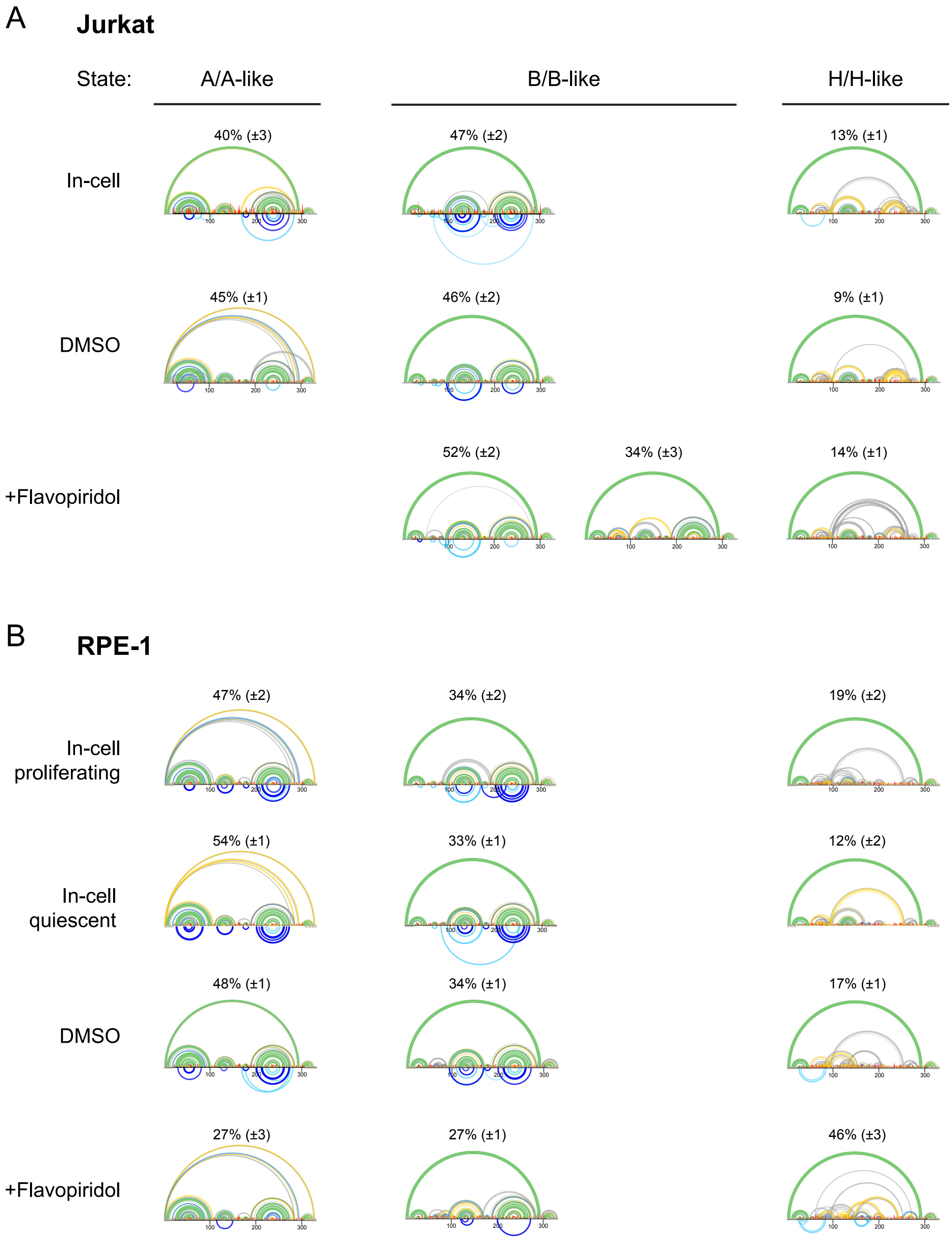
A
Jurkat
State:
A/A-like
B/B-like
H/H-like
In-cell
40% (±3)
47% (±2)
13% (±1)
DMSO
45% (±1)
46% (±2)
9% (±1)
+Flavopiridol
52% (±2)
34% (±3)
14% (±1)
B
RPE-1
In-cell proliferating
47% (±2)
34% (±2)
19% (±2)
In-cell quiescent
54% (±1)
33% (±1)
12% (±2)
DMSO
48% (±1)
34% (±1)
17% (±1)
+Flavopiridol
27% (±3)
27% (±1)
46% (±3)
